# Episodic memory formation in unrestricted viewing

**DOI:** 10.1101/2022.05.24.485821

**Authors:** Andrey R. Nikolaev, Inês Bramão, Roger Johansson, Mikael Johansson

## Abstract

The brain systems of episodic memory and oculomotor control are tightly linked, suggesting a crucial role of eye movements in memory. But little is known about the neural mechanisms of memory formation across eye movements in unrestricted viewing behavior. Here, we leverage simultaneous eye tracking and EEG recording to examine episodic memory formation in free viewing. Participants memorized multi-element events while their EEG and eye movements were concurrently recorded. Each event comprised elements from three categories (face, object, place), with two exemplars from each category, in different locations on the screen. A subsequent associative memory test assessed participants’ memory for the between-category associations that specified each event. We used a deconvolution approach to overcome the problem of overlapping EEG responses to sequential saccades in free viewing. Brain activity was time-locked to the fixation onsets, and we examined EEG power in the theta and alpha frequency bands, the putative oscillatory correlates of episodic encoding mechanisms. Three modulations of fixation-related EEG predicted high subsequent memory performance: 1) theta increase at fixations after *between-category* gaze transitions, 2) theta and alpha increase at fixations after *within-element* gaze transitions, 3) alpha decrease at fixations after *between-exemplar* gaze transitions. Thus, event encoding with unrestricted viewing behavior was characterized by three neural mechanisms, manifested in fixation-locked theta and alpha EEG activity that rapidly turned on and off during the unfolding eye movement sequences. These three distinct neural mechanisms may be the essential building blocks that subserve the buildup of coherent episodic memories during unrestricted viewing behavior.

## 1 Introduction

Life events consist of visual, semantic, contextual, and other associated features stored as coherent episodic memory representations. Episodic memory allows us to mentally travel in time to relive past events with considerable detail and specificity (Tulving, 1983). A critical aspect when building such memory representations is that the visual field of our perception is spatially limited. At any given time, we can only apprehend information in full acuity in a small spot, determined by the size of the fovea. We overcome this limitation by constantly shifting our visual focus with eye movements. In effect, these visual “samples” of the world constitute the pieces we successively bind together into meaningful, coherent episodic memories (e.g., Ryan et al., 2020; Voss et al., 2017). Nonetheless, in the neuroscience of human memory, memory formation has primarily been studied in experimental paradigms where the study material is presented in a single location on the screen and where eye movements are typically treated as artifacts. Thus, little is currently known about the neural mechanisms of episodic memory formation across eye movements. In the present study, we set out to capture the neural signatures of building coherent episodic memories via eye movements at the level of individual gaze fixations. To this end, we apply a state-of-the-art approach of simultaneous eye-tracking and EEG recording and analysis in a free-viewing memory task.

Episodic memory formation critically hinges upon binding together the different elements that constitute an event. Sequential saccadic eye movements organize these relationships as a mechanism that links different aspects of the environment into coherent memory representations (for overviews, see Ryan et al., 2020; Voss et al., 2017). Eye movements also serve an important role during memory retrieval (Damiano and Walther, 2019; Johansson and Johansson, 2014; Johansson et al., 2022; Wynn et al., 2020), and a longstanding theory holds that the stored representations include the sequences of eye movements established during memory formation, which supposedly carry information about linked event elements (Brandt and Stark, 1997; Noton and Stark, 1971).

The clock frequency of this ‘linking’ is the theta rhythm (4-7 Hz). We typically make about 3-4 saccades in natural viewing per second. Such visual sampling frequency corresponds to the theta rhythm (Amit et al., 2017; Otero-Millan et al., 2008). The saccadic theta rhythm is part of a periodic mechanism of attentional deployment (Hogendoorn, 2016). It also corresponds to theta oscillations of the hippocampus, which is of crucial importance for associative memory formation (Cohen and Eichenbaum, 1993). The hippocampus also has tight anatomical connections with brain structures that control oculomotor behavior (Pierrot-Deseilligny et al., 2004; Shen et al., 2016). Recent data suggest that hippocampal theta oscillations support saccade guidance during memory formation on a fixation-by-fixation basis (Jutras et al., 2013; Kota et al., 2020; Kragel et al., 2021; Kragel et al., 2020). In this way, the theta cycle may organize sequential visual inputs and enables the binding of event elements into a coherent memory representation (Herweg et al., 2020).

The relevance of theta oscillations for memory processes has long been established in studies using stimulus-response paradigms with fixed stimulus presentations. For instance, increased frontal theta power during episodic memory encoding predicts subsequent retrieval performance (Hsieh and Ranganath, 2014; Klimesch, 1996). The increase in theta power, i.e., theta synchronization, is often accompanied by a decrease in alpha (8-13 Hz) power, i.e., alpha desynchronization (Hanslmayr et al., 2016; Waldhauser et al., 2012). Alpha desynchronization may represent the fidelity of memory representations for the constituent elements of episodes (Hanslmayr et al., 2016). Thus, alpha and theta oscillations serve essential but distinct functions of episodic memory formation: representing elements (alpha) and binding them together into a coherent whole (theta). It is currently unknown how these mechanisms engage during the sequential sampling of event elements across saccades to allow later episodic remembering of the event as a whole. In the current study, we track how these neural mechanisms subserve the buildup of episodic memories during unrestricted viewing behavior.

Brain activity and eye movements must be examined simultaneously to allow insight into neural mechanisms involved in unrestricted viewing. Over the last decade, the combination of EEG and eye-tracking has become a powerful tool for studying perceptual and cognitive processes in unrestricted viewing (Coco et al., 2020; Devillez et al., 2015; Dimigen et al., 2011; Fudali-Czyz et al., 2018; Körner et al., 2014; Tyson-Carr et al., 2020), including attention and memory across saccades. For instance, presaccadic EEG activity has been found to predict successful working memory of task-relevant scene elements in a change blindness task (Nikolaev et al., 2011). However, despite the apparent benefit of utilizing EEG-eye movement coregistration, attempts to apply it are virtually absent in the episodic memory literature. One explanation may be the vast methodological challenges when combining these techniques, including the effects of overlapping saccades on EEG activity in free viewing (Dimigen and Ehinger, 2021; Dimigen et al., 2011; Nikolaev et al., 2016). Moreover, low-level oculomotor effects of eye movement characteristics can systematically differ between experimental conditions and distort the EEG, creating differences that are likely to be confounded with the effects of the experimental conditions. To overcome these and other challenges as described below, the present study applied a regression-based deconvolution approach that recovers unknown isolated neural responses given the measured EEG and the latencies of experimental events (Dimigen and Ehinger, 2021; Ehinger and Dimigen, 2019). This approach ultimately allowed us to isolate and track the neural correlates of episodic memory formation across eye movements.

In the present study, we investigate how theta and alpha activity subserve the binding of separate event elements into a coherent episodic memory across eye movements. To this end, we introduced a free-viewing episodic memory task in which participants encoded multiple events. Each event was defined by six distinct event elements (images), which always comprised two exemplars from each of three categories (faces, places, objects) presented in different locations on the screen. The two exemplars from the same category always appeared close together, forming a pair, whereas the distance between categories (i.e., between the pairs) was much greater (Fig. 1). Participants were asked to memorize the images that appeared together on the same screen, i.e., belonged to the same event. In a subsequent associative memory task, we tested participants’ episodic memory for the between-category associations (i.e., associations between elements from different categories) within each event (Fig. 1).

**Fig. 1.**
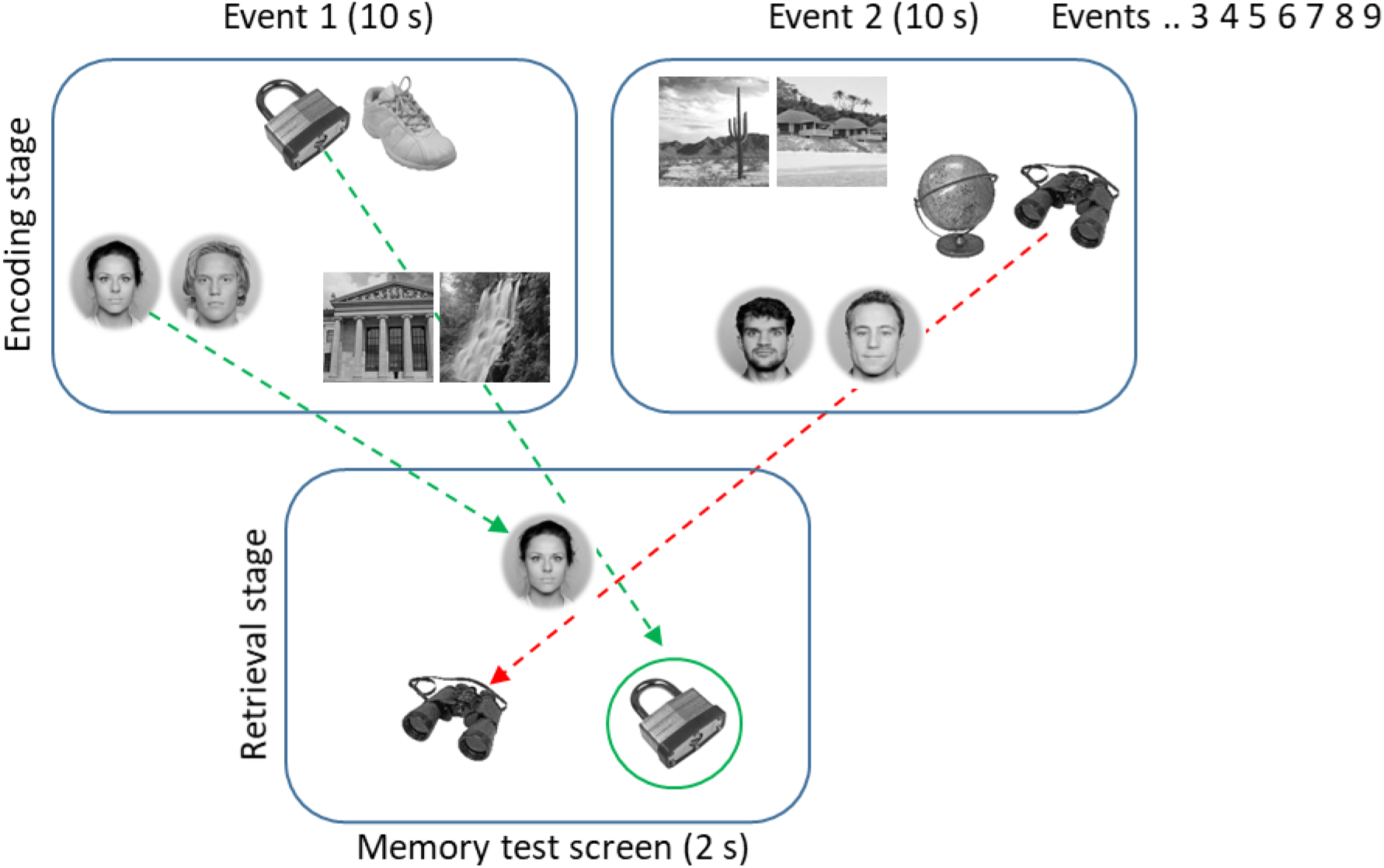
At encoding, nine events were sequentially presented. Each event comprised two exemplar images from three different categories (faces, places, objects) and was presented for 10 s. After a 30-s distractor task, episodic memory was assessed by testing all between-category associations from each event. The test screen (presented for 2 s) included a cue image at the top with two images at the bottom from a different category. Participants were asked to indicate which of the two images had appeared together with the cue in the previously encountered events. In this example, the correct response is the lock to the right, as it appeared together with the face cue in Event 1, whereas the binoculars to the left appeared in Event 2. Note that the angular distances between categories were much larger on the actual encoding screen.

To form a coherent event representation during encoding, participants critically need to engage in sequential eye movements, where each event element gets visually attended to on one or multiple occasions. Thus, to successfully establish the element associations specifying each event, we expected the encoding process to critically depend on participants’ gaze transitions between the elements (cf. Kragel et al., 2021), as well as on the visual sampling of each individual element (cf. Liu et al., 2017; Loftus, 1972; Olsen et al., 2016). As the ensuing associative memory task concerned between-category associations, we expected gaze transitions between categories to be of particular importance and, thus, more predictive of memory performance than transitions between exemplars from the same category (associations that participants were not asked to retrieve). Based on previous research by Kragel et al. (2021), we also expected repeated cross-category gaze transitions to facilitate successful event formation.

We simultaneously recorded EEG and eye movements during event encoding to examine theta and alpha activity related to such gaze behaviors during memory formation. In our analytical approach, we first sought to verify that the oscillatory nature of the EEG signals coregistered with eye movements in our free-viewing paradigm corresponds to the encoding-related neural signatures typically found in comparable studies with fixed stimulus presentation (Hanslmayr et al., 2016). To achieve the central goals of the present study, we then corrected EEG from overlapping effects of eye movements using a deconvolution approach (Ehinger and Dimigen, 2019) and extracted the EEG power relative to fixation onset. Since this was the first study to examine scalp-recorded EEG coregistered with eye movements during a free-viewing associative memory task, we first investigated the time course of fixation-related EEG during the whole 10-s interval of event encoding. Then, we examined EEG power related to three types of fixations: (1) fixations succeeding a *between-category* saccade (i.e., a saccade moving from one element to another across categories); (2) fixations succeeding a *between-exemplar* saccade (i.e., a saccade moving from one element to another within the same category); and (3) fixations succeeding a *within-element* saccade (i.e., a saccade moving from one point to another within the same element). In general, we expect all saccades between event elements to be important for building a coherent episodic representation. However, as our memory task required the retrieval of between-category associations, we reasoned that saccades between categories should be vital and predictive of subsequent performance, in contrast to between-exemplar saccades supporting associations that were not called for. Thus, by analyzing both saccade types, we can examine the specificity of gaze transitions between event elements upon subsequent memory performance as a function of task relevance. This approach allowed us to examine alpha and theta activity related to functionally distinct gaze behaviors contributing with complementary aspects to the episodic memory representation: eye movements that serve to (1) sample visual information from the individual event elements and crucially (2) link the different event elements into a coherent whole.

## 2 Methods

### 2.1 Participants

Thirty-six healthy adults participated in the experiment in exchange for a gift card in a shopping mall (approx. €10). Three participants were excluded because of system crashes during the data collection and another three due to below-chance memory performance (the chance level threshold was set at 53.2%, which was obtained from the permutation procedure indicating chance performance in a two-alternative forced choice (2AFC) task (‘left’ or ‘right’ responses in a memory test) for the 5% confidence level and for a total number of 648 test trials). Finally, two participants were excluded due to ceiling performance (> 90%), i.e., too few incorrect trials for the planned subsequent memory analysis. The final group included 28 participants (20 females; mean age 23.3; age range 18-32).

The study was conducted in accordance with the Swedish Act concerning the Ethical Review of Research involving humans (2003:460). All participants gave written informed consent, and the study followed the local ethical guidelines at Lund University. Moreover, the study followed the Lund University safety protocol for conducting EEG, eye-tracking, and behavioral experiments in the context of the COVID-19 pandemic.

### 2.2 Stimuli

A total of 324 photo images from three different categories: faces, places, and objects, were used. Faces were selected from the Oslo Face Database (Chelnokova et al., 2014), which provided consent for the publication of the faces that were used in the figures. Objects were selected from the Bank of Standardized Stimuli (Brodeur et al., 2010). Places were selected from the Michael Tarr lab scene database (http://www.tarrlab.org/). The images were converted to black-and-white, and each image’s pixel intensity was normalized to the average pixel intensity of all images. In the present study, multiple simultaneously presented images defined an ‘event’. Each event comprised three image pairs (two exemplars from each category) distributed across distinct locations on the display (Fig. 1). Two exemplars of each a category were necessary for sufficient task complexity with respect to the target-distractor dynamics (i.e., exemplars from the same category both within and across events). Moreover, this particular configuration was found to give optimal memory performance (60-70% correct) for nine events in the block (see below) in pilot behavioral studies, where we manipulated the number of exemplars (one or three) per category together with the number of events per block. Exemplars from the same category were close to each other, whereas the different categories were more separated. Specifically, each image had a square shape with a side of 4.2 degrees of visual angle (dva). The width of a pair (two exemplars from one category) was 8.5 dva, and the distance between the centers of the three pairs was 17.9 dva. The three pairs were placed around the perimeter of an invisible circle with a radius of 9.2 dva centered at the center of the screen. The location of the pairs on the circle was random for each event, but the pairs were always separated by arcs of 120 degrees. The distribution of images and their pairing across events was random for each participant. The images were presented on a grey background.

### 2.3 Procedure

Participants sat in a dimly lit room, with their heads stabilized with a chin rest. Stimuli were presented at a viewing distance of 62 cm on an EIZO FlexScan EV2451 monitor with a resolution of 1920 × 1080 pixels and a refresh rate 60 Hz, which was part of the Tobii Pro Spectrum eye tracker.

The experiment comprised six blocks with the possibility to take a short break between each block. Each block included two main stages: memory encoding and retrieval, with a distractor task in between (Fig. 1). Participants self-initiated each block by pressing the space bar. During encoding, nine events were presented for 10 s each. Before each event, a fixation cross was presented for a random interval between 1 and 1.5 s. Participants were instructed to memorize the faces, places, and objects that appeared together in the same event. The encoding stage was followed by a distractor task, where participants were asked to count backward in steps of 7 from a random 3-digit number for 30 s. Pilot experiments indicated that the selected number of exemplars, events, and blocks gave rise to an optimal memory performance (60-70% correct).

A cued recall task followed three seconds after the distractor task, which assessed episodic memory by testing retrieval of the between-category associations of each event. A test screen with three images, one at the top and two at the bottom, was presented for 2 s (Fig. 1). Each test screen was preceded by a fixation cross randomly jittered between 1 and 1.5 s. The image at the top served as the retrieval cue. Participants were asked to indicate which of the two images at the bottom had appeared together with the cue during encoding, i.e., to indicate the target. The distractor was an exemplar from the same category as the target but selected from a different event studied no more than two events before or after the target event. Participants responded by pressing the left or right arrow keys of a standard keyboard. Participants were thereafter asked to indicate how confident they were in their response. The question “Are you sure?” was displayed on the screen, and participants responded: “Sure”, “Maybe”, or “Guess”, by pressing the left, down, and right arrow keys, respectively.

Between-category associations were tested for each image in each event on four trials. For example, a face image was tested twice with place images (once as cue and once as target), and twice with object images (once as cue and once as target). The cue-target status of categories was counterbalanced across blocks, such that in the odd blocks, the cue-target status was faces-places, places-objects, objects-faces, and in the even blocks, the corresponding cue-target status was places-faces, objects-places, faces-objects. Thus, memory for each event was assessed by tests of 12 different associations. In total, 108 associations were tested in each block (12 × 9 = 108 tests for the nine events within a block) with randomized order of presentation. Participants were not explicitly instructed that only the between-category associations were tested but could infer that from the exhaustive practice sessions (described below).

Memory performance for the associates may depend on varying encoding difficulty for the three categories. We eliminated such potential confound by the counterbalancing of individual exemplars and their screen location across events, blocks, and participants, during both encoding and testing screens (Fig. 1). We also evaluated performance as a function of the type of cross-category association within an event: faces – places, faces – objects, places – objects (reported in 3.1).

Prior to the main experiment, two practice sessions with increasing difficulty (three events in the first session, six events in the second session) were conducted in which participants received feedback on their performance after each test trial. The gradual increase in difficulty over the two practice sessions served to prepare participants for what they could expect in the main experiment (where nine events per block were presented). Ten associative memory tests were delivered in the practice sessions, in which between-category associations were evenly distributed over events and categories. Images in the practice sessions were not repeated in the main experiment and were not repeated over the two practice sessions. The overall duration of the experiment was about two hours. The experiment was programmed in PsychoPy v.2020.2.4 (Peirce et al., 2019).

### 2.4 Simultaneous EEG and eye movement recording

EEG and eye movements were recorded concurrently throughout the experiment. EEG data were recorded using a Grael 4K EEG amplifier (Neuroscan, Compumedics Limited, Australia) at a sampling frequency of 2048 Hz using 31 Ag/AgCl electrodes positioned according to the extended 10-20 International system (Easycap GmbH, Germany). Two electrodes were mounted over the left and right mastoids, using the latter as the recording reference. The ground electrode was AFz. Additional electrodes recorded the vertical and horizontal electrooculograms (VEOG and HEOG): the VEOG electrodes were placed above and below the right eye, the HEOG electrodes were placed at the left and right outer canthi of two eyes. Electrode impedances were kept below 5 kΩ.

The Tobii Pro Spectrum eye tracker (Tobii, Stockholm, Sweden) recorded movements of both eyes with the sampling frequency of 600 Hz. 9-point calibration and validation routines were conducted before the first block. An error of 1 dva between calibration and validation was tolerated: if it was larger, the calibration was repeated. The eye tracking was controlled from the PsychoPy experimental program via the open-source toolbox Titta (Niehorster et al., 2020). The experimental program ran on the same computer that controlled the eye tracker. To synchronize stimulus presentation and EEG recording, a transistor-transistor logic (TTL) signal was sent via a parallel port from the stimulus presentation computer to the EEG system at the beginning and end of each block, encoding event, and memory test.

### 2.5 Data analysis

Only data from the encoding stage were analyzed. Analysis was conducted with custom scripts for R (Team, 2020) as well as for MATLAB R2020a (The MathWorks Inc., Natick, Massachusetts), using the EEGLAB toolbox (Delorme and Makeig, 2004), the EYE-EEG extension (Dimigen et al., 2011) for EEGLAB, and the Unfold toolbox (Ehinger and Dimigen, 2019).

#### 2.5.1 Behavioral analysis

Memory performance for each event was assessed by retrieval accuracy weighted by confidence, such that correct responses were scored as 3 for “Sure” responses, as 2 for “Maybe” responses, and as 1 for “Guess” responses. Each image from each event occurred either as a cue or a target in four memory tests trials. Thus, each image can in total receive a score ranging from 0 (incorrect responses in all four tests) to 12 (correct responses accompanied by four “sure” responses in all four tests). To calculate a memory score for the whole event, we then summed the scores for the six elements of each event (ranging from 0 to 72). For the EEG analysis, events were divided into “high memory” and “low memory,” according to the subsequent memory scores. For each participant, high and low memory was calculated by a median split of that participant’s memory scores for the whole events.

#### 2.5.2 Eye movement analysis

Fixations and saccades were detected from participants’ right eye using the velocity-based algorithm for saccade detection (Engbert and Mergenthaler, 2006) implemented in the EYE-EEG extension. Fixations were considered located on an image if within 0.5 dva of the outer contour of the image. Since participants cannot adequately perform a memory test if they do not visually process (fixate) all images while event encoding, we excluded from all analyses the events where fixations were detected only on two of the three categories (on average 3±4 (mean ± SD) events per participant). Such events occurred because of eye-tracking issues mainly towards the end of the experiment (see the Supplementary materials and Fig. S1 for details). Fixations outside the images on different parts of the display were assigned to an ‘other’ saccade type, which was used as a reference level in the deconvolution modeling (2.5.5).

Saccades were classified according to whether they occurred between categories (between-category saccades), between the two exemplars of a category (between-exemplar saccades), and within an exemplar (within-element saccades). In addition, we distinguished between the first visit to an image and revisits to it. Altogether, we defined five different types of saccades (Fig. 3A):

1. Between-category saccades: first visits
2. Between-category saccades: revisits
3. Between-exemplar saccades: first visits
4. Between-exemplar saccades: revisits
5. Within-element saccades

We expected these saccade types to differentially contribute to forming a coherent event representation. Specifically, in our associative memory task, we tested episodic memory for between-category associations (i.e., associations between exemplars from different categories) within each event. The formation of these associations supposedly depends on the corresponding between-category saccades during encoding. In contrast, associations established by between-exemplar saccades (i.e., gaze transitions between associations incidental to the task) were not tested and did not expect to predict subsequent memory performance. Thus, between-category and between-exemplar saccades support associations between event elements that were or were not critical to the success of the subsequent memory test, respectively.

Revisits, i.e., refixations of previously visited locations, have proven to be of particular interest in memory-related gaze behavior. Specifically, refixations can increase the sampling of information necessary for successful memory formation (Mirza et al., 2016; Turk-Browne, 2019; Voss et al., 2011) and recover information lost or missed during scanning (Gilchrist and Harvey, 2000; Körner and Gilchrist, 2008; Meghanathan et al., 2019; Tatler et al., 2005). Our previous coregistration studies have shown that refixations differ in the allocation of attention during saccade planning from ordinary fixations to locations visited once (Nikolaev et al., 2018) and that refixation planning is goal-dependent (Meghanathan et al., 2020). Moreover, we found that information acquisition at fixations to locations that are revisited later is distinct when compared to the other fixations (Nikolaev et al., 2022). Refixations may also play an important role in the formation of episodic memories. A recent study combining eye-tracking with intracranial EEG (Kragel et al., 2021) demonstrated that presaccadic hippocampal theta activity predicts whether an upcoming saccade will be a first visit or a revisit. Based on these previous findings, we expected EEG activity during revisits to be predictive of the successful binding of the event elements.

Additionally, we examined scrutinizing saccades within an exemplar (*within-element*), as it is well-known that the cumulative number of fixations on a visual stimulus predicts subsequent memory of that stimulus (Liu et al., 2017; Loftus, 1972; Olsen et al., 2016). This allowed us to examine two types of memory mechanisms: the first related to eye movements that bind the elements into a coherent event, and the second related to eye movements that sample visual information from individual elements.

#### 2.5.3 EEG preprocessing

The goal of our study required fixation-related analysis of EEG in short epochs time-locked to fixation onsets in the encoding interval. This analysis used deconvolution modeling to correct for the overlapping effects of saccades on EEG. Deconvolution modeling involves time regression, which requires continuous EEG; thus, preprocessing was performed on continuous EEG. In addition, we analyzed EEG time-locked to the event onset during the entire 10-s encoding interval. The preprocessing for this analysis was the same as for the fixation-related analysis. We used the following preprocessing and deconvolution pipelines developed in our previous research (Nikolaev et al., 2022).

We down-sampled EEG signals to 256 Hz. The EEG and eye-tracking data were synchronized using the function *pop_importeyetracker* from the EYE-EEG extension. Saccades, fixations, event onsets, and bad eye-tracking intervals were inserted into the EEGLAB data structure.

The preprocessing pipeline included a series of cleaning functions from EEGLAB. The *pop_cleanline* function removed power line noise using multi-tapering and a Thompson F-statistic. The *clean_artifacts* function removed flat-line channels, low-frequency drifts, noisy channels, and short-time bursts. This function is based on artifact subspace reconstruction (ASR), which compares the structure of the artifactual EEG to that of known artifact-free reference data (Kothe and Jung, 2016). The tradeoff between artifactual and retaining brain activities depends on the ASR parameter, which we set to 20 according to the recommendations by Chang and colleagues (Chang et al., 2020).

Ocular artifacts were removed with the *OPTICAT* function (Dimigen, 2020). First, EEG was high-pass filtered at 2 Hz to suppress large deviations from baseline due to summation of the corneo-retinal artifacts during sequential eye movements. Next, 30-ms segments around saccade onsets were obtained and re-appended to EEG to ‘overweight’ the contribution of the saccadic spike activity in the EEG input to independent component analysis (ICA). Then, the ICA weights obtained after ICA training on these filtered data were transferred to the unfiltered version of the same data. Finally, the ratio between the mean variance of independent components during saccade and fixation intervals was calculated. If the ratio was higher than 1.1, the corresponding independent components were considered saccade-related and removed (Plöchl et al., 2012).

The automatic classifier (*pop_iclabel*) with the following settings (the probability ranges of ICs to be an artifact and excluded are given in brackets): Brain [0 0], Muscle [0.4 1], Eye [0.9 1], Heart [0.05 1], Line Noise [0.4 1], Channel Noise [0.4 1], Other [0.4 1] revealed independent components related to remaining artifacts (Pion-Tonachini et al., 2019). On average, 16.9 (SD = 3.3, range 10-25) components per participant were removed. EEG was re-referenced to average reference. The removed channels (mean = 1.1, SD = 1 per participant) were interpolated with spherical spline interpolation.

Theta and alpha power were extracted by applying the procedure from (Ossandón et al., 2020). We filtered the continuous EEG with a low cut-off of 4 Hz and high cut-off of 7 for the theta band and with a low cut-off of 8 Hz and high cut-off of 13 for the alpha band using the *pop_eegfiltnew* function with default settings. We Hilbert-transformed the filtered signal and obtained the instantaneous power. We normalized the EEG power at each sampling point to its ratio with the respective channel mean power in a baseline interval. In the analysis of the encoding interval, the baseline interval was set from -200 to 0 ms before the onset of the corresponding event. In the fixation-related analysis, to avoid an interval of saccade execution just before fixation onset, the baseline interval was set from -200 to -100 ms before the onset of the corresponding fixation interval. The power values were log-transformed.

#### 2.5.4 Oscillation detection

To test the presence of theta and alpha oscillations in the encoding interval we used the approach for detection of transient oscillatory bursts, which separates periodic (oscillatory) from aperiodic activity. This approach is based on comparison of the observed power spectrum with the aperiodic background 1/f activity. It has been implemented in a number of methods, such as better oscillation detection (BOSC) (Caplan et al., 2001) and its further developments, such as extended BOSC (eBOSC) (Kosciessa et al., 2020) and the fBOSC (Seymour et al., 2022), which includes the ‘fitting oscillations and one over f’ (FOOOF) algorithm (Donoghue et al., 2020). We searched for theta and alpha oscillations in the 10-s encoding interval using the fBOSC toolbox for MATLAB (Seymour et al., 2022). The fBOSC method was particularly suitable for our goals because it assumes that the 1/f fit is not necessarily linear across all frequencies in log-log space, and allowed us to fit an aperiodic ‘knee’ in the power spectrum. The presence of such a ‘knee’ is especially likely in data with pronounced theta rhythmicity (Seymour et al., 2022), which is quite expected in our data.

To detect oscillations, fBOSC computes two thresholds: a power threshold estimated from the aperiodic background spectrum, which we set at the 95^th^ percentile of the theoretical probability (chi-squared) distribution of power, and a duration threshold, which we set at 3 oscillatory cycles. Since oscillations may distort the background spectrum, fBOSC excludes oscillatory peaks from its calculation. To do this, fBOSC performs an initial 1/f fit, then iteratively models oscillatory peaks above this background and removes them from the spectrum. Next, fBOSC models the 1/f power spectrum as nonlinear by using a variable knee parameter (see Seymour et al. (2022) for details on its estimation). We applied fBOSC to single EEG trials of the 10-s encoding interval. The results of the single-trail modeling were averaged across all participants for low and high memory events separately. To analyze the frequency distribution of the detected oscillations and their relationship to memory, we used ‘abundance’, which is considered the most informative metric in burst analysis (Kosciessa et al., 2020). Abundance is the duration of a rhythmic episode relative to the length of the analyzed segment, which can range from 0 to 1, where 0 means no burst and 1 means a single continuous burst.

#### 2.5.5 Deconvolution

To overcome the problems associated with EEG-eye movement coregistration in unrestricted viewing behavior described above, we used deconvolution modeling implemented in the Unfold toolbox (Ehinger and Dimigen, 2019). The toolbox allowed us to correct overlapping EEG responses to sequential saccades in free viewing, as well as to account for several possible detrimental covariates. The deconvolution modeling is based on multiple regression, as traditional mass-univariate ERP modeling, which has been used for post-hoc control of multiple simultaneous discrete and continuous covariates (Smith and Kutas, 2015). However, mass-univariate modeling requires a prespecified analysis window. Therefore, when applied to correct the overlapping effects of unrestricted eye movements on EEG, it cannot account for the varying temporal overlap caused by variable fixation durations between adjacent eye movements (Dimigen and Ehinger, 2021). Deconvolution corrects for overlapping effects with variable fixation intervals by time-expansion of the design matrix in the duration of the entire (continuous) EEG recording (see Fig. 4 in Dimigen and Ehinger (2021) for an illustration of time-expansion). Furthermore, the correction method implemented in the Unfold toolbox accounts for the nonlinear effects of oculomotor covariates (e.g., saccade size) on EEG by the combination of linear deconvolution with nonlinear spline regression, as used in the generalized additive model (GAM) (Wood, 2017). The resulting time series of regression coefficients, or betas, represent non-overlapping brain responses for each predictor and correspond to subject-level ERP waveforms in traditional ERP analysis. Thus, the outputs of the toolbox are beta coefficients, which represent partial effects for the predictors of interest, adjusted for the covariates included in the model.

These covariates included the low-level oculomotor effects of eye movement characteristics and the ordering factors in time courses of an encoding trial, a block, and an entire experiment. Specifically, we considered the low-level oculomotor characteristics of fixation duration and saccade size and direction, which are known to affect the fixation-related EEG (Dimigen et al., 2011; Nikolaev et al., 2016). Furthermore, we considered three ordering factors. Since the time course of a trial with multiple eye movements may affect the fixation-related EEG activity (Fischer et al., 2013; Kamienkowski et al., 2018), we considered the rank of a fixation in relation to all other fixations that occurred during the entire encoding interval of an event. The rank was assigned to each fixation as an ordinal number of this fixation in the encoding interval. Behavioral results (see below) revealed a nonlinear effect of event order within a block and a strong linear effect of block order on subsequent memory. We accounted for these factors by including them in the deconvolution model.

The Unfold toolbox involved the following procedures, which were sequentially applied to the continuous EEG of each participant.

To account for the experimental conditions of interest and possible confounding factors in the same model, we used the following Wilkinson notation of the model formula:

~~~
Fixation: y ∼ 1 + cat(MemoryPerformance) + rank + iBlock + spl(iEventPerBlock,4) +
spl(duration,5) + spl(sac_amplitude,5) + circspl(sac_angle,5,-180,180)
Event: y ∼ 1
Levels of *MemoryPerformance*: ‘other’ (reference level), ‘low memory’, ‘high memory’
~~~

This formula includes the multiple effects at fixations, as well as the onset of the event screen. Specifically, for fixations, the formula considers the fixation onset (y ∼ 1, i.e., the intercept), a categorical predictor of the subsequent memory effect (‘MemoryPerformance’) and continuous predictors: fixation rank (‘rank’), a block order in an experiment (‘iBlock’), an event order in a block (‘iEventPerBlock’), fixation duration (‘duration’), size (‘sac_amplitude’) and direction (‘sac_angle’) of the preceding saccade. In this example formula, the ‘MemoryPerformance’ predictor includes three levels: ‘other’ fixations (see Eye movement analysis above), ‘low memory’ and ‘high memory’ (see Behavioral analysis above). We used treatment (dummy) coding with ‘other’ fixations as the reference level (intercept) relatively to which other levels (i.e., ‘low memory’ and ‘high memory’) of the ‘MemoryPerformance’ predictor were estimated. The inclusion of ‘other’ fixations allowed us to account for the overlapping effects on EEG produced at the latencies of ‘other’ eye movements. To effectively account and control for these overlapping effects, we need to input in the model all fixations that occurred in the experiment, without exception. Thus, the “other” fixations, together with the fixations assigned to low and high memory levels, were essential levels of the ‘MemoryPerformance’ predictor. However, because the “other” fixations cannot explain memory for the event, they were never used in further comparisons. Since fixation duration, and saccade size and direction have nonlinear effects on EEG (Dimigen and Ehinger, 2021; Nikolaev et al., 2016), we modeled them with a basis set of five spline predictors (circular splines were used to model saccade direction). Since we found that the event order in the block has a nonlinear effect on the memory score (Fig. 2B), and we assumed its nonlinear effect on the EEG, we modeled it also with spline predictors. Spline knots were placed on the percentiles of the covariates for each participant. For events, we considered only their onsets (y ∼ 1), i.e., the intercept that described the overall shape of EEG evoked by the event screens. In further analyses, we modified the ‘MemoryPerformance’ predictor in the formula by inserting predictor levels depending on the analysis goals, as specified below in each section of Results. All other predictors were constant in all analyses.

**Fig. 2.**
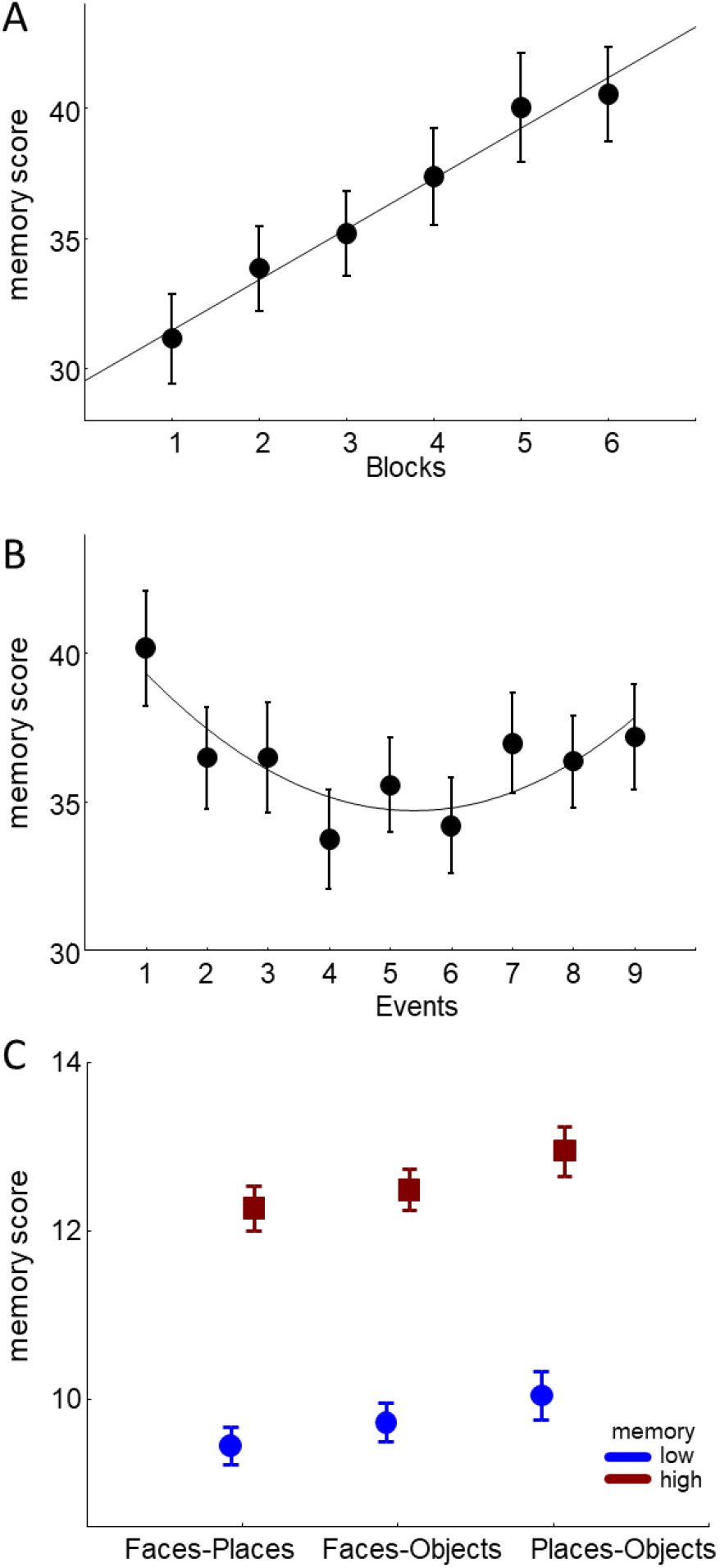
Memory performance weighted by confidence in the time course of an experiment (A) and a block (B). The thin lines indicate significant linear (A) and quadratic (B) fits. (C) Memory performance weighted by confidence for paired associations between particular categories within an event and divided according to low and high memory performance in the entire event (as in the EEG analysis).

To recover isolated EEG responses (betas) to each fixation and to each event screen best explaining the continuous EEG, we created a design matrix (Fig. S3A) and time-expanded it in a window between −200 and +400 ms around fixation and event onsets. This length of the time window was motivated by the need to keep the number of iterations for fitting the deconvolution model before model convergence reasonably small (e.g., <400) to avoid overfitting. The number of iterations depends on the number (and quality: spline or not) of predictors and the number of sampling points. At the pilot stage of the project, our experimentation with the window length using the selected predictors showed that the length of -200+400 ms is optimal. Time expansion involved duplicating and shifting the predictors of the mass univariate linear regression design matrix for each time lag.

The time-expanded design matrix (Fig. S3B) spanned the duration of the entire EEG recording. As a result, instead of estimating multiple linear models for all time points, we estimated one linear model, simultaneously estimating all fixation and event betas. The variable temporal distance between sequential fixations allowed the separation of the overlapping effects.

Before fitting the model, we excluded irrelevant intervals from modeling by filling them with zeros in the time-expanded design matrix. Filling with zeros preserved the timing of continuous EEG. If we would remove these intervals completely, we might remove along with them the eye movements that produce overlapping effects on clean EEG parts used in the analyses of interest. Specifically, these intervals corresponded to the distractor task, memory tests, intervals between events, breaks between blocks, and bad eye-tracking intervals, leaving only intervals of free viewing exploration of event screens during encoding. We fitted the deconvolution model to each of the 31 electrodes using the iterative Least Squares Minimal Residual algorithm for sparse design matrices (Fong and Saunders, 2011).

The raw beta coefficients obtained for categorical predictors of interest after fitting the deconvolution model represent the “pure” effects of the predictors, which could be already interpreted and compared statistically. The effects could be understood as a rough analog of difference waves relative to the reference level of the categorical predictor. Consequently, the curves of the “pure” effects have highly wiggly shapes, similar to the difference waves, oscillating around zero, as illustrated in Fig. S4. This makes it difficult to compare deconvolution results with the waveforms in the traditional EEG literature. To solve this problem, we need to add the model intercept onto the difference wave. However, in the case of nonlinear modeling with splines, calculating the intercept is not straightforward as it would be in the case of linear models. Specifically, in nonlinear modeling with splines, the classical model-intercept is ill-defined, since there is no clear reference level for the nonlinear basis set (the set which transforms predictor values by weighting beta coefficients). To resolve this issue, we estimated all nonlinear effects at their respective mean predictor levels and added these estimates to the beta coefficients of the categorical predictors. For example, if we aim to adjust our memory effect to saccade amplitude, and the mean saccade amplitude for a given participant is 7 dva, we evaluated the nonlinear effect at 7 dva and added the resulting value to the beta coefficients of the memory effect. This procedure ensures that all levels of the memory effect are visualized at the same estimated saccade amplitude of 7 dva. The described solutions are implemented in the functions *uf_predictContinuous* and *uf_addmarginal* of the Unfold toolbox, which we applied to the raw beta coefficients for each participant. The resulting waveforms then became comparable to the participant-level averages in traditional EEG analysis.

#### 2.5.6 Statistical analysis

Unless indicated otherwise, we used a repeated measures ANOVA for the statistical analysis. To handle violation of sphericity, we applied the Huynh-Feldt correction for p-values associated with two or more degrees of freedom. Tukey HSD test was used for post-hoc analyses. The statistical analyses were performed with STATISTICA 10 (StatSoft Inc., Tulsa, OK, USA) and R (R Core Team, 2020).

Data analysis involved three major stages. First, we aimed to examine whether the theta and alpha activity obtained from EEG signals coregistered with eye movements in unrestricted viewing would yield results consistent with the previous episodic memory studies with fixed stimulus presentation (Hanslmayr et al., 2016). Since theta and alpha activity represent distinct functions of episodic memory formation and have complex relationships, we analyzed them in separate ANOVA models. We segmented theta and alpha EEG power before deconvolution into 10-s trials relative to the onsets of the encoding screens (events). We compared the mean EEG power in five 2-s time bins of 10-s trials between events with low and high subsequent memory performance. Furthermore, we detected the presence of theta and alpha oscillations in the encoding interval with the fBOSC method (Seymour et al., 2022). We considered the duration (‘abundance’) of theta and alpha oscillations for low and high subsequent memory. In these and all subsequent EEG analyses, to capture effects with potentially different topographical distributions, we grouped electrodes in eight regions of interest (ROIs). Specifically, we selected three electrodes around the landmark electrodes of the International 10-20 System of Electrode Placement: F3, F4, C3, C4, P3, P4, O1, O2 (the inset in Fig. 4). These groups of electrodes constituted eight ROIs over frontal, central, parietal and occipital brain regions of the left and right hemisphere (FL, FR, CL, CR, PL, PR, OL, OR). For each ROI, we averaged power waveforms across three electrodes since regional averaging of electrodes provides a more reliable estimate of activity in a region than a single measurement (Dien and Santuzzi, 2005). Eight ROIs were symmetrically about the sagittal axis and systematically distributed over the scalp. Thus, these analyses allowed us to evaluate memory-related changes of theta and alpha power over the entire encoding interval.

Second, we evaluated the general characteristics of fixation-related power extracted from the deconvolved EEG. We estimated the theta and alpha power at the first peak of their cycle about 100 ms after the fixation onset (see 4.3 for justification of such selection). We averaged the power values 40 ms before and after the peak, that is, between 60 and 140 ms after the fixation onset. To evaluate the time course of fixation-related EEG power during the 10-s encoding interval, we divided EEG power related to successive eye movements into three bins according to fixation rank. We compared memory-related changes of theta and alpha power across rank bins in eight ROIs. Thus, this analysis was analogous at the fixation level to the first analysis at the event level.

The third analysis was devoted to the main goal of our study – to compare fixation-related EEG *after* the five types of saccades described in section 2.5.2. We performed three separate analyses on each saccade type because not all conditions were matched for all saccade subtypes. Specifically, we could distinguish between First visits and Revisits for Between-category and Between-exemplar saccades but not for Within-element saccades. All analyses compared the fixation-related theta and alpha power between low and high memory for the same time window as in the second analysis and eight ROIs. This analysis provided a detailed picture of associations during the formation of coherent episodic memory at the fixation level. To emphasize findings directly relevant to the goals of our study, we report effects and interactions in the Results section only when they involved the factors of memory performance (High vs. Low) and gaze transition status (First visits vs. Revisit). Complete statistical results, including effects and interactions between ROIs and hemispheres, are reported in Table S2 in Supplementary materials.

## 3 Results

### 3.1 Behavioral results

Average memory performance (regardless of confidence) across 28 participants was 68.6 ± 8.33 % (mean ± SD). The memory score weighted by confidence for the low memory condition was 27.8 +/- 6.8 and for the high memory condition 48.8 +/- 9 (mean +/- SD). Memory performance changed in the time course of a block and experiment (Fig. 2). We evaluated these changes with a repeated measures ANOVA on the memory scores weighted by confidence with the factors of Event (nine levels) and Block (six levels). There were significant main effects of Event (F(8, 216) = 3.7, p = .004, ε = .62) and Block (F(5, 135) = 9.8, p < .001, ε = .95)), but no interaction between them. To characterize the effects of Event and Block, we used planned comparisons with the linear and quadratic contrasts. The contrast analysis revealed significant effects for the linear contrast for Block (F(1, 27) = 31.2, p < .001) and for the quadratic contrast for Event (F(1, 27) = 16.8, p < .001), but not for the linear contrast for Event (F(1, 27) = 0.7, p = .4). These results indicate a prominent learning effect during the experiment, as well as possible primacy and recency effects in the time course of a block. To evaluate the contribution of confidence to memory scores, we repeated the above analyses on memory scores that were not weighted by confidence. The results were qualitatively the same (Supplementary materials and Fig. S2), indicating that confidence contributes little to our measure of memory performance.

To compare encoding of different categories, we computed three separate memory scores (weighted by confidence), considering only paired associations within an event: faces – places, faces – objects, places – objects. Then we divided these memory scores according to low and high memory performance in the entire event (as in the EEG analysis). We applied a repeated-measures ANOVA on the memory scores of category associations with factors of Memory (Low vs. High) and Association type (faces – places, faces – objects, places – objects). We found effects of Memory (F(1, 27) = 375, p < .001) and Association type (F(2, 54) = 10.5, p = .001, ε = .66), and no interaction (F(2, 54) = 0.5, p = .63) (Fig. 2C). The post-hoc test showed similar results for low and high memory performance: no difference in memory scores between associations of faces – places and faces – objects (both p > .06) and significant differences in memory scores between associations of faces – places and places – objects (both p < .001), as well as between associations of faces – objects and places – objects (both p < .021). These results statistically confirmed the low recall of faces reported by most participants in post-experiment interviews. More importantly, differences in performance across categories were equally distributed between the low and high memory events used as conditions in the EEG analyses and, therefore, could not bias the EEG results.

### 3.2 Eye movement results

The dynamics of eye movement behavior during the 10-s encoding interval is illustrated in Fig. 3B by the fixation rank probability density for each type of preceding saccade. At first, participants tended to visit all categories quickly with between-category saccades. Then, between-exemplar saccades became predominant. The number of within-element saccades was relatively stable throughout the whole encoding interval. Towards the latter part of the interval, between-category revisits became more frequent.

To determine if, and to what degree, the saccade categories predicted subsequent memory performance, we analyzed the relationship between the cumulative number of saccades (in each category) within an event and subsequent memory performance for that event. The between-category saccades for first visits were not analyzed since there were always 3 per event. We computed the number of the remaining four saccade types for each event and pooled these numbers with the corresponding event memory scores over 28 participants. Since the number of saccades for different types varied quite widely (Fig. 3C), we could not establish a single categorical predictor of the number of saccades for all saccade types to include them in the same ANOVA design. Therefore, we applied a one-way ANOVA on the event memory score with the number of the saccades as the predictor for each saccade type separately (bins with less than 20 values at the left and right ends of the distributions were excluded). The results showed that memory for the event significantly increased with the cumulative number of between-category revisits (F(8, 1392) = 3.8, p < .001) and with the cumulative number of within-element saccades (F(14, 1360) = 3.1, p < .001) (Fig. 3C).

**Fig. 3.**
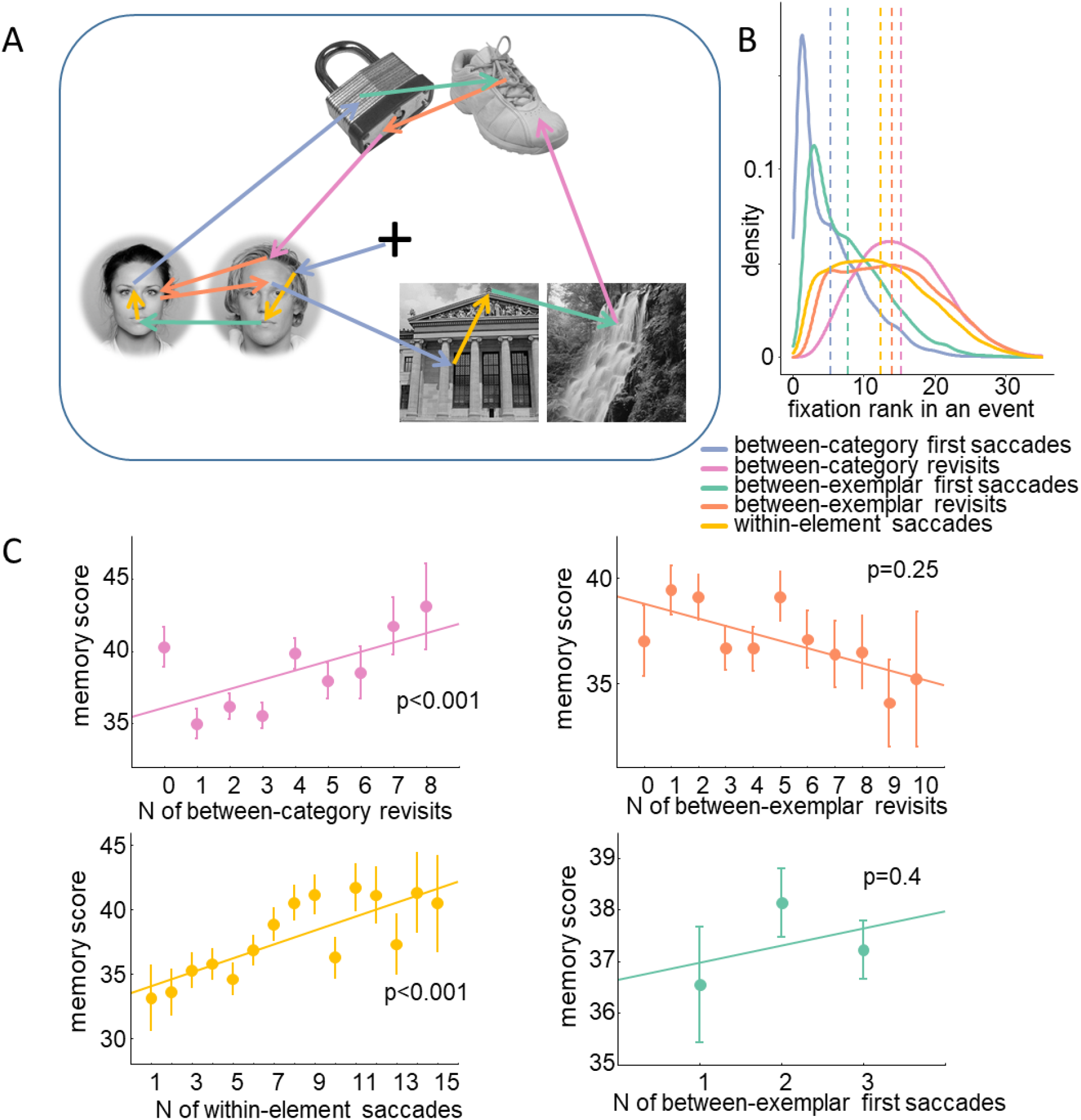
A: an example scanpath illustrating five types of saccades. B: Probability density estimation of the fixation rank in a 10-s event interval pooled across 28 participants. The vertical lines indicate mean ranks of the corresponding fixation type. C: Dependence between the occurrence frequency of saccade types and the memory score. Significance of the dependence obtained with one-way ANOVA is indicated in each panel. Error bars indicate standard errors of means across 28 participants.

To be successful in the retrieval task, association from all three categories need to be considered during event encoding. Two different gaze patterns can achieve this: (1) engaging in active gaze transitions between each of the three categories, or (2) engaging in active gaze transitions between only two of the categories, i.e., when the association between the first visited category and the last visited category is inferred without an active gaze transition between the categories. In the present data set, gaze transitions involving only two categories characterized approximately one-third of the events. These events were distributed between participants approximately normally (the Lilliefors test could not reject a null hypothesis of normality of the distribution: d = 0.12, p > .2). To evaluate whether the two eye movement patterns are associated with different subsequent memory performance, we compared event memory scores between these pattern types with a paired t-test. We found no difference in memory between the two patterns (t(27) = 0.05, p = .96). The memory score and the number of events involving only two categories did not correlate across participants (r = .25, p = .2). These findings suggest that event memory does not require gaze transitions between all three categories.

We also compared the rate of eye movements between low and high subsequent memory performance. The eye movement rate normalized to the total number of saccades for each participant was much higher in the low (.53, .025) than high (.47, .025) (mean, SD per participant) memory condition (t(27) = 5.7, p < .001).

In sum, the eye movement results show that between-category saccades (revisits) and within-element saccades predict subsequent memory performance, whereas between-exemplar saccades do not. The associative memory task determined these dependencies, which required strong associations between the categories, whereas associations between the exemplars of a category were less important to the task. Within-element saccades were necessary for acquiring sufficient visual information about each image, a prerequisite for establishing a strong representation of each element, and for recognizing them during retrieval.

### 3.3 EEG results at the event level

#### 3.3.1 Subsequent memory effects at the event level

To assess how theta and alpha power at encoding predicts subsequent memory at the time scale of the entire encoding interval, we segmented the EEG power (after removing all artifacts but before deconvolution) from -0.2 to 10 s relative to the event onset and baseline corrected it to the interval from – 0.2 to 0 s before the event onset (Fig. 4A, B). We smoothed the individual power waveforms with a Savitzky-Golay filter, using filter order 3 and filter length 201 sampling points (R package ‘signal’). Events were split between low and high memory performance (the number of low memory events was 28.5 (1.35); the number of high memory events was 25.1 (1.74) (mean (SD) per participant). To investigate the modulation across a 10-s encoding interval, we created five time-bins of 2 s each and averaged the power values in these bins (Fig. 4C, D). We applied a repeated-measures ANOVA on the power values with the factors of Memory (Low vs. High), ROI (Frontal, Central, Parietal, Occipital), Hemisphere (Left vs. Right), and Time (5 bins). For theta power, we found an interaction between Memory and Time (F(4, 108) = 4.9, p = .001, ε = .99). Post-hoc test revealed that this interaction occurred due to higher power for the high than low memory events for the 4^th^ and 5^th^ time bins (all p < .02), i.e., between 6 and 10 s at the end of the encoding interval (Fig. 4C).

**Fig. 4.**
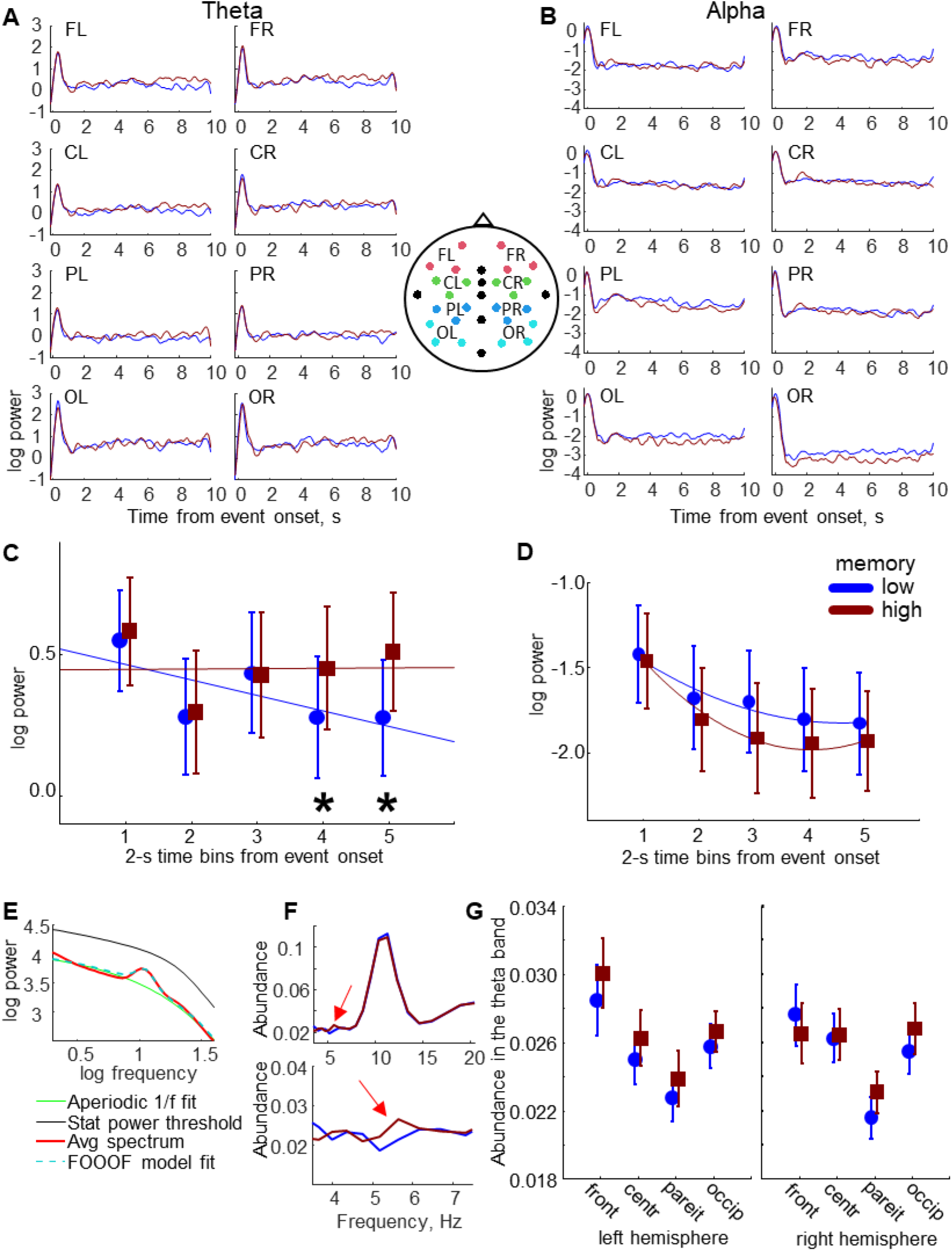
Theta (A) and alpha (B) power time-locked to the event onset during a 10-s encoding interval for low and high subsequent memory performance. The power is grand averaged across 28 participants in 8 ROIs selected as indicated in the inset between panels A and B. (C – D) The time course of the grand-averaged theta (C) and alpha (D) power waveforms and the mean-error plots (mean +/- SEM) for five 2-s time bins. Significant results of the post-hoc test for the interaction between Memory and Time bins are indicated with asterisks. In (C) the straight lines indicate a linear fit, whereas in (D) the curved lines indicate a quadratic fit, best reflecting the power changes over time in the respective bands. (E) Log-log plot of the EEG power spectrum averaged across 28 participants over the left parietal area. (F – G) The duration of the rhythmic episode relative to the length of the analyzed segment (‘abundance’) averaged across 28 participants over the left parietal area for low and high memory performance. (F) (top) Abundance for the frequency range 3-20 Hz. (bottom) Abundance in the theta band (close-up of the plot above). The arrows indicate the abundance increase for high memory performance. (G) The mean-error plots (mean +/- SEM) of abundance in the theta band for 8 ROIs.

For alpha power, there was a clear trend towards decreased power in high-memory versus low-memory events, most noticeably over the occipital areas (Fig. 4B, D), which, however, did not reach significance. The change of alpha power over time appears to be nonlinear. To test the properties of this change statistically, we applied linear and quadratic contrasts to the time bins as planned comparisons. For this analysis, we used only the data from the occipital ROIs, where the memory effect on alpha power was most prominent (Fig. 4B). We found that the quadratic fit more adequately describes the memory effect on the alpha activity over time, revealing a significant difference between high and low memory events (F(1, 27) = 5.0, p = .03), than the linear fit, which did not reveal a significant effect between high and low memory events (F(1, 27) = 0.5, p = .47). For completeness, we also applied the linear and quadratic contrasts to the time bins for theta power (all ROIs). Here the results were the opposite: there was a significant difference between low and high memory events for the linear fit (F(1, 27) = 17.5, p < .001) but not for the quadratic fit (F(1, 27) = 2.8, p = .1).

In sum, EEG analysis of the entire encoding interval revealed the expected neural signatures, where higher subsequent memory performance was associated with increased theta synchronization and alpha desynchronization during encoding (Hanslmayr et al., 2016). Consistent with the idea of a gradual buildup of an episodic memory representation, theta increase was more prominent towards the end of the encoding interval. Theta increase was accompanied by alpha decrease, whose relationship with subsequent memory performance over time was nonlinear and best described by a quadratic fit.

#### 3.3.2 Detection of theta and alpha oscillations at the event level

The finding that subsequent memory performance is associated with theta and alpha activity raises the question of whether memory-related changes in spectral pattern result from neural oscillations pertinent to encoding (Hanslmayr et al., 2016) or instead reflect increases in broadband activity or shifts in spectral tilt. We detected theta and alpha oscillations using the fBOSC method, which considered the possible nonlinearity of the 1/f power spectrum (Seymour et al., 2022). Fig. 4E shows the averaged power spectrum, the aperiodic 1/f component, and the FOOOF model fit for the left parietal area for high memory events (the results for all ROIs and both events are shown in Fig. S5A). The grand-averaged results were about the same for low- and high-memory events. The power spectrum had a clear peak in the alpha range and a deviation from the aperiodic fit at low frequencies.

Although the grand-averaged power did not reach the power threshold, multiple oscillation episodes were detected at the single-trial level. We compared the duration of these oscillations (‘abundance’) between low and high memory. Fig. 4F shows the abundance averaged across participants for the left parietal area for the frequency range 3-20 Hz (top) and for the close-up in the theta range (bottom). The abundance for all ROIs is shown in Fig. S5B. The arrows point to the frequency interval of about 5-6 Hz, where the abundance was larger for the high than low memory events. The abundance at the peak alpha frequency was visually larger for the low than high memory events. To test these observations statistically, we averaged the abundance within the theta and alpha bands for each participant and analyzed it using a repeated-measures ANOVA with the factors of Memory (low vs. high), ROI (frontal, central, parietal, occipital), and Hemisphere (left vs. right). For the theta band, we found main effects of Memory (F(1, 27) = 4.8, p = .038) and a tendency for an interaction between Memory, ROI, and Hemisphere (F(3, 81) = 2.3, p = .086, ε = .98). Fig. 4G suggests that this interaction tendency may occur because of the increased abundance over the parietal regions for high memory events. For the alpha band, there was no Memory effect (F(1, 27) = 0.4, p = .56).

Thus, the abundance analysis shows that a long duration of theta oscillations throughout the encoding interval predicts high memory performance. The duration of alpha oscillations is much longer than the duration of theta oscillations, as indicated by the prominent increase of abundance in the alpha band (Fig. 4F). But the duration of alpha oscillations throughout the encoding interval is not predictive of memory performance. This, however, does not exclude their involvement in memory processes at the fixation level.

### 3.4 EEG results at the fixation level

Since the deconvolution framework is a novel approach to free-viewing behavior in tasks of episodic memory formation, we first evaluated how it changes the power waveforms (see Supplementary materials and Fig. S6 for details).

#### 3.4.1 Subsequent memory effects at the fixation level

Before analyzing the data for the different saccade categories, we wanted to assess if the EEG findings at the event level could be reproduced at the fixation level. Thus, we tested the subsequent memory effect on fixation-related theta and alpha power by contrasting the fixation-related EEG overall eye movements within an event. We divided the memory scores into three bins according to low, medium, and high memory events. We divided the fixations into three bins according to fixation rank. In the formula for deconvolution modeling, we used ten levels of categorical predictors: all possible combinations of three memory and three rank bins and ‘other’. The number of fixations used in the deconvolution modeling of each memory and rank bin condition is presented in Table S1.

We applied a repeated-measures ANOVA on the EEG power averaged in the window from 60 to 140 ms after the fixation onset with the factors of Memory (low vs. high, we excluded the medium memory bin to increase the memory contrast), Rank (3 bins), ROI (frontal, central, parietal, occipital) and Hemisphere (left vs. right). For theta power, we found interactions between Memory and Rank (F(2, 54) = 3.6, p = .038, ε = .96) (Fig. 5B) and between Memory and ROI (F(3, 81) = 3.6, p = .023, ε = .86) (Fig. 5C). Post-hoc tests revealed that the Memory-Rank interaction occurred due to higher power for the high than low memory in rank bin 3 (p = .02). Fig. 5A shows power waveforms for this rank bin. The Memory-ROI interaction occurred due to higher power for the high than low memory over the frontal (p = .006) and central (p = .001) areas. The difference maps indicate that this effect was largest in rank bin 3 (Fig. 5D).

**Fig. 5.**
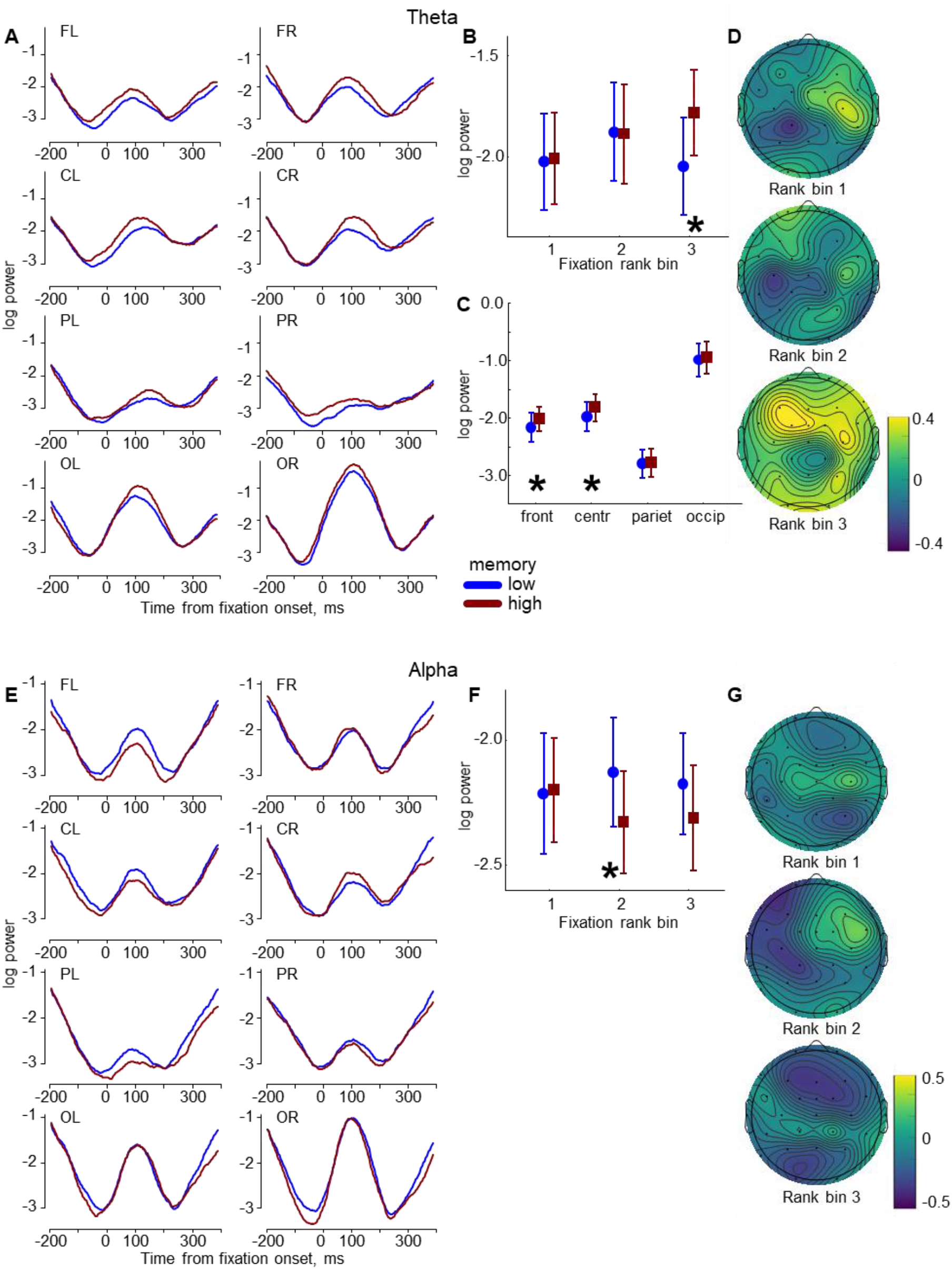
Theta and alpha fixation-related grand-averaged (N=28) power in the analysis of three fixation rank bins. (A) Theta power in 8 ROIs for rank bin 3. (B – C) The mean-error (mean +/- SEM) plots of the theta power in the interval from 60 to 140 ms after the fixation onset. (B) Memory-Rank interaction. (C) Memory-ROI interaction. (D) The theta power difference maps: high minus low memory. (E) Alpha power in 8 ROIs for rank bin 2. (F) The Memory-Rank interaction over the left hemisphere. (G) The alpha power difference maps: high minus low memory conditions. Significant results of post-hoc tests are indicated with asterisks.

For alpha power, we found an interaction between Memory, Rank, and Hemisphere (F(2, 54) = 5.0, p = .01, ε = .98) (Fig. 5E-G). Post-hoc tests revealed that the interaction occurred due to a power decrease for the high than low memory in rank bin 2 (Fig. 5F) over the left hemisphere (p = .002); the similar trend in rank bin 3 did not reach post-hoc significance (p = .14). There was also an interaction between Memory, Rank, ROI, and Hemisphere (F(6, 162) = 2.4, p = .04, ε = .84), but the post-hoc tests showed no significant differences.

To summarize, the fixation-related EEG results corroborate and extend what was found at the event level, i.e., theta increase and alpha decrease during encoding covary with subsequent memory performance. These effects were also more prominent toward the end of the encoding interval. Next, we analyzed EEG power at fixations following the five saccade types of interest.

#### 3.4.2 Subsequent memory effects across saccade types

We focused on EEG power in the fixation intervals succeeding the five saccade types (Fig. 3A). For this purpose, we specified ten levels of categorical predictors in the deconvolution model formula: the five saccade types x memory performance (low, high). The number of fixations that were used in the deconvolution modeling of each saccade and memory condition is presented in Table 1. “Other” fixations were the reference level, as in the previous model.

**Table 1.** The number of fixations used in the deconvolution modeling of each saccade type per participant: mean (range, SD) across 28 participants.

|  | low memory | high memory |
| --- | --- | --- |
| Between-category saccades: first visits | 83 (57-96, 8.6) | 69 (57-78, 7.1) |
| Between-category saccades: revisits | 82 (28-158, 35.4) | 74 (18-156, 32.6) |
| Between-exemplar saccades: first visits | 64 (39-83, 12.3) | 55 (37-75, 11.9) |
| Between-exemplar saccades: revisits | 115 (47-214, 51.7) | 95 (31-177, 44.1) |
| Within-element saccades | 187 (101-321, 67.4) | 164 (88-288, 60.5) |

##### 3.4.2.1 Between-category saccades

Here, we examined EEG power in the fixation intervals succeeding the between-category saccades that we predicted to support tested associations between categories (Fig. 6A). We applied a repeated measures ANOVA on fixation-related theta and alpha power with the factors of Gaze transition status (First visits vs. Revisit), Memory (Low vs. High), ROI (Frontal, Central, Parietal, Occipital), and Hemisphere (Left vs. Right). For theta power, we found a significant main effect of Gaze: theta power was higher during first visits than revisits (F(1, 27) = 5.0, p = .03). We also found an interaction between Gaze and ROI (F(3, 81) = 4.2, p = .01, ε = .94). More importantly, we found an interaction between Gaze, Memory, and Hemisphere (F(1, 27) = 11.8, p = .002) (Fig. 6B-E). Post-hoc tests revealed that this interaction occurred due to higher power for the high than low memory for revisits over the left hemisphere (p < .001) (Fig. 6E). For alpha power, no main effects of Memory (F(1, 27) = 3.1, p = .09) or Gaze (F(1,27) = 1.7, p = .2) emerged (nor an interaction between them, F(1, 27) = 0.5, p = .5).

**Fig. 6.**
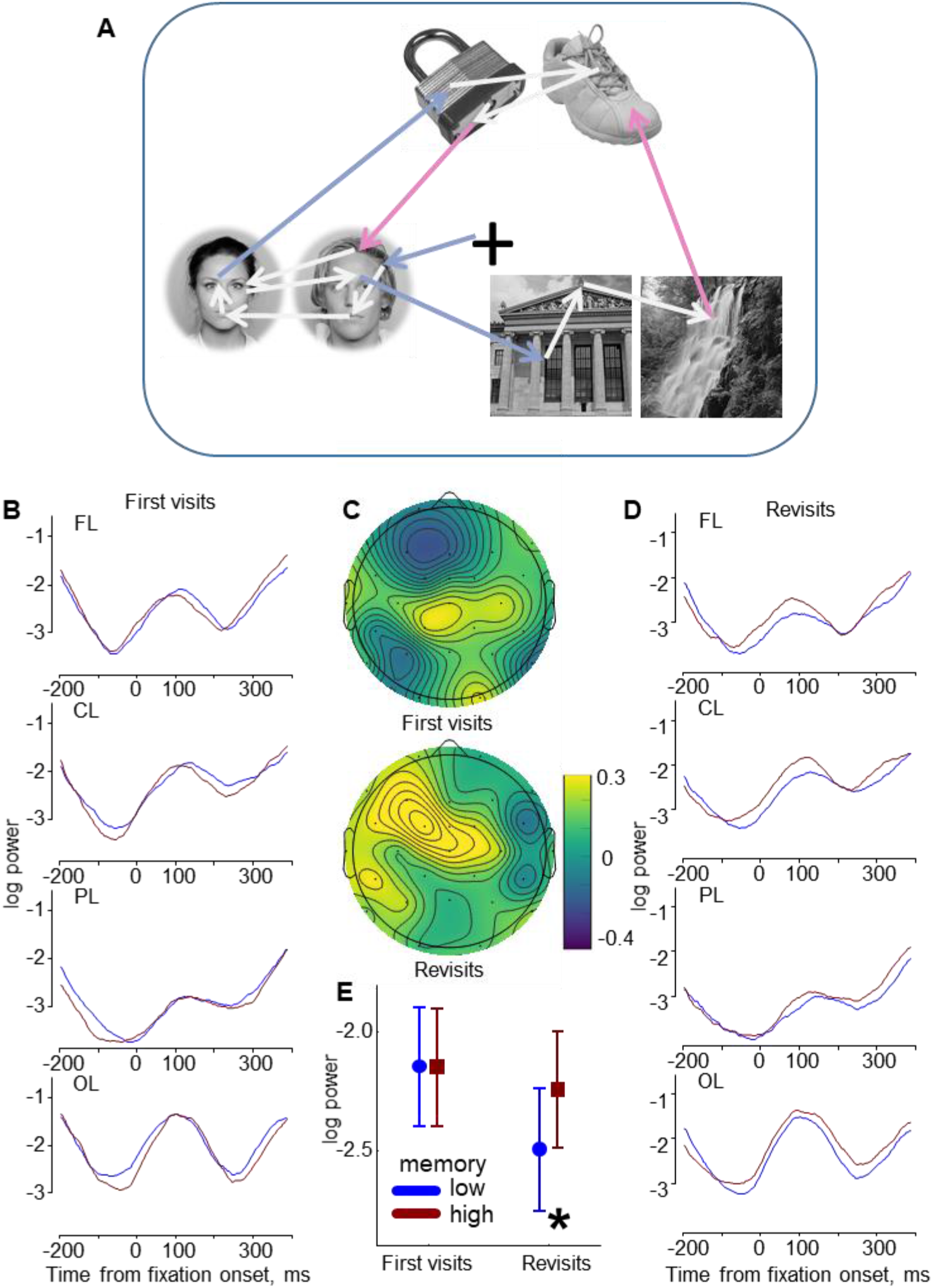
Fixation-related EEG after between-category saccades. (A) An example scanpath illustrating a subset of between-category (colored) saccades preceding the fixation intervals used in the EEG analysis. The grey arrows are saccades not included in this analysis. (B, D) The grand-averaged (N=28) theta power over the left hemisphere for first visits (B) and revisits (D). (C) The power difference maps: high minus low memory conditions. (E) The mean-error (mean +/- SEM) plot of the power over the left hemisphere in the interval from 60 to 140 ms after the fixation onset for first visits and revisits. Significant results of post-hoc tests are indicated with an asterisk.

Thus, theta power increase over the left hemisphere after between-category revisits predicts subsequent event memory performance. This theta memory effect is consistent with the eye-movement analyses, where subsequent memory performance improved with the cumulative number of between-category revisits during encoding (Fig. 3C).

##### 3.4.2.2 Between-exemplar saccades

Next, we examined EEG power in the fixation intervals succeeding between-exemplar saccades, i.e., gaze transitions establishing element associations that were not tested in the subsequent memory test (Fig. 7A). As in the previous analysis, we used a repeated measures ANOVA with a 2×2 design: Gaze transition status (First visit vs. Revisit) x Memory (Low vs. High), ROI (Frontal, Central, Parietal, Occipital), and Hemisphere (Left vs. Right). For alpha power, we found the main effects of Memory (Fig. 7B-D), where power was lower for high than low memory performance (F(1,27) = 5.1, p = .03). For theta power, no significant effects of Memory (F(1,27) = 0.3, p = .6) or Gaze (F(1,27) = 2.1, p = .16) emerged (nor an interaction between them F(1,27) = 1.5, p = .24).

**Fig. 7.**
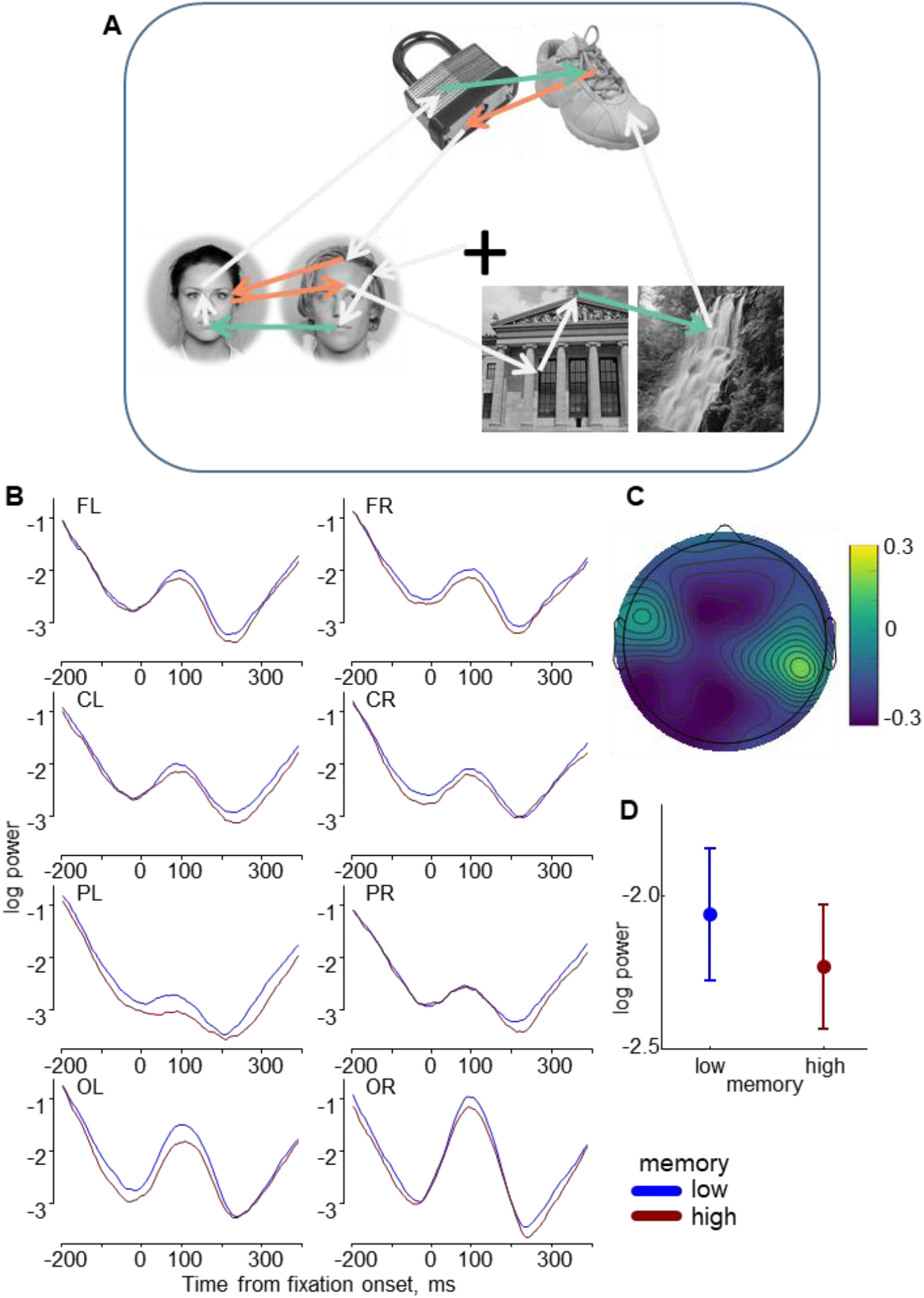
Fixation-related EEG after between-exemplar saccades. (A) An example scanpath illustrating a subset of between-exemplar (colored) saccades preceding the fixation intervals used in the EEG analysis. The grey arrows are saccades not included in this analysis. (B) The grand-averaged (N=28) alpha power for fixations after between-exemplar saccades. (C) The difference alpha power maps: high minus low memory conditions. (D) The mean-error (mean +/- SEM) plot of the alpha power in the interval from 60 to 140 ms after the fixation onset.

As expected, theta power after between-exemplar saccades did not predict subsequent event memory performance for the tested between-category associations. This is consistent with the eye-movement result, where the cumulative number of between-exemplar saccades did not influence subsequent memory performance (Fig. 3C). The effect of lower alpha power for high memory was unexpected, but may reflect other mechanisms, apart from binding *per se*, that are important for optimal memory formation.

##### 3.4.2.3 Within-element saccades

Finally, we analyzed theta and alpha power in the fixation intervals after within-element saccades (Fig. 8A). For this saccade type, we did not distinguish between first visits and revisits, so the ANOVA design included three factors: Memory (Low vs. High), ROI (Frontal, Central, Parietal, Occipital), and Hemisphere (Left vs. Right). For theta power, we found an interaction between Memory and ROI (F(3, 81) = 3.4, p = .025, ε = .95) (Fig. 8B, C). Post-hoc tests revealed that this interaction occurred due to higher theta power for high than low memory over the frontal (p = .002) and central (p = .03) areas (Fig. 8D). For alpha power, we found an interaction between Memory and Hemisphere (F(1, 27) = 4.9, p = .035) (Fig. 8E-F). Post-hoc tests revealed that this interaction occurred due to higher alpha power for high than low memory over the right hemisphere (p = .01) (Fig. 8G).

**Fig. 8.**
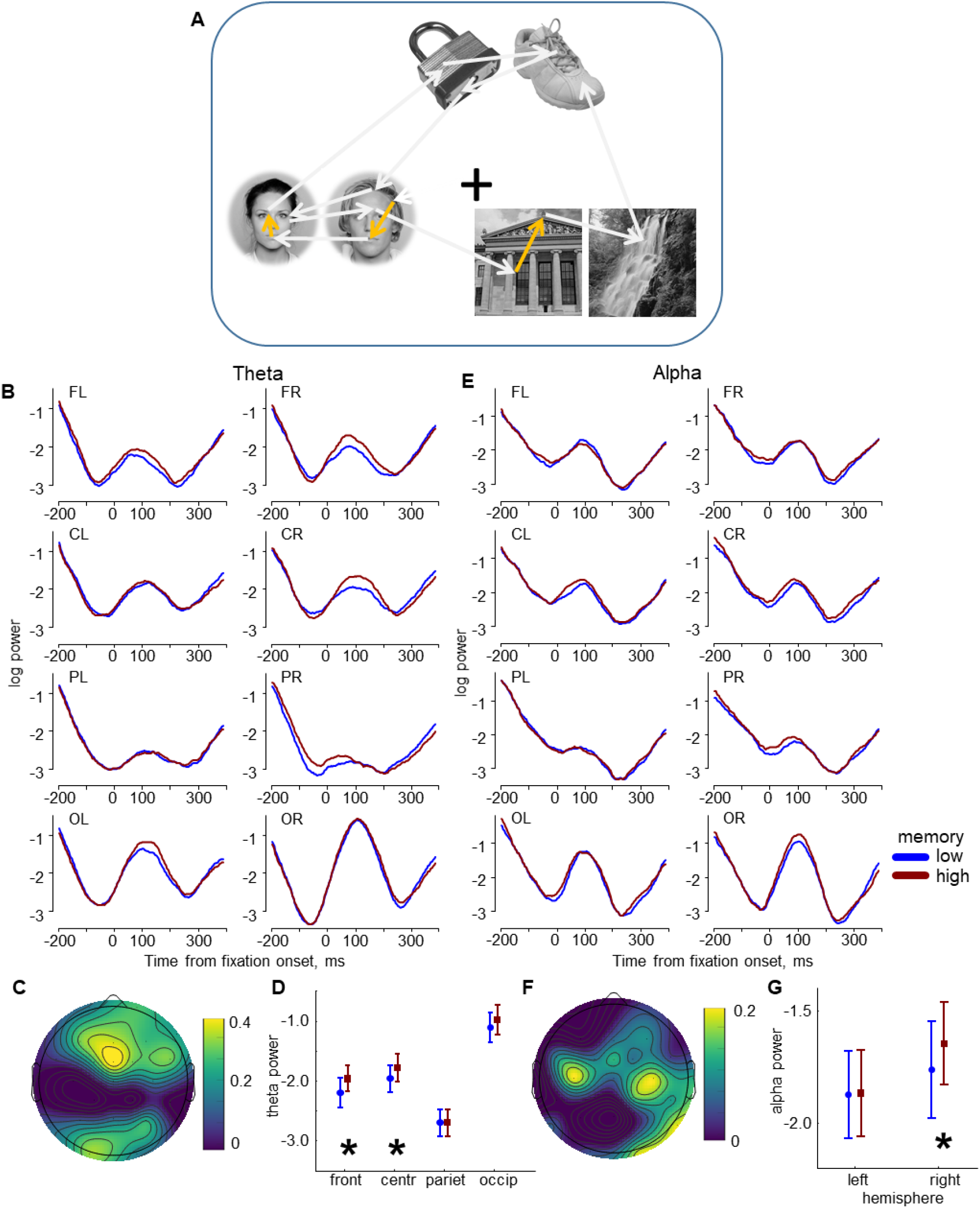
Fixation-related EEG after within-element saccades. (A) An example scanpath illustrating a subset of within-element (colored) saccades preceding the fixation intervals used in the EEG analysis. The grey arrows are saccades not included in this analysis. The grand-averaged (N=28) theta (B) and alpha (E) power for 8 ROIs. The theta (C) and alpha (F) power difference maps: high minus low memory conditions. (D, G) The mean-error (mean +/- SEM) plot of the power in the interval from 60 to 140 ms after the fixation onset. (D) Memory-ROI interaction for theta power. (G) Memory-Hemisphere interaction for alpha power. Significant results of post-hoc tests are indicated with asterisks.

Thus, the theta power related to within-element saccades predicts subsequent memory performance similarly as for the between-category revisits. Moreover, for both saccade types, subsequent memory increased as a function of the cumulative number of saccades during encoding (Fig. 3C). Importantly, however, the topographies of the theta memory effects for the two saccade types were *distinct*: a left central maximum for between-category revisits (Fig. 6C) and a frontal maximum for within-element saccades (Fig. 8C), suggesting dissociable neural mechanisms.

To test the observed topographical dissociation of the two theta effects, we extracted the ‘within-element effect’ as the difference between theta power in high and low memory performance for fixations after within-element saccades. We extracted the ‘between-category effect’ as the difference between theta power in high and low memory performance for fixations after between-category revisits. A repeated-measures ANOVA on the theta power difference with factors of Theta effect (‘within-element’ vs. ‘between-category’), ROI (Frontal, Central, Parietal, Occipital), and Hemisphere (Left vs. Right) revealed an interaction between Theta effect and Hemisphere (F(1, 27) = 9.0, p = .006). The post-hoc test revealed higher power for the between-category effect over the left hemisphere (p = .03) (Fig. 9). This finding indicates topographical differences for the two theta effects and suggests that they are associated with distinct neural mechanisms operating in the same frequency band.

**Fig. 9.**
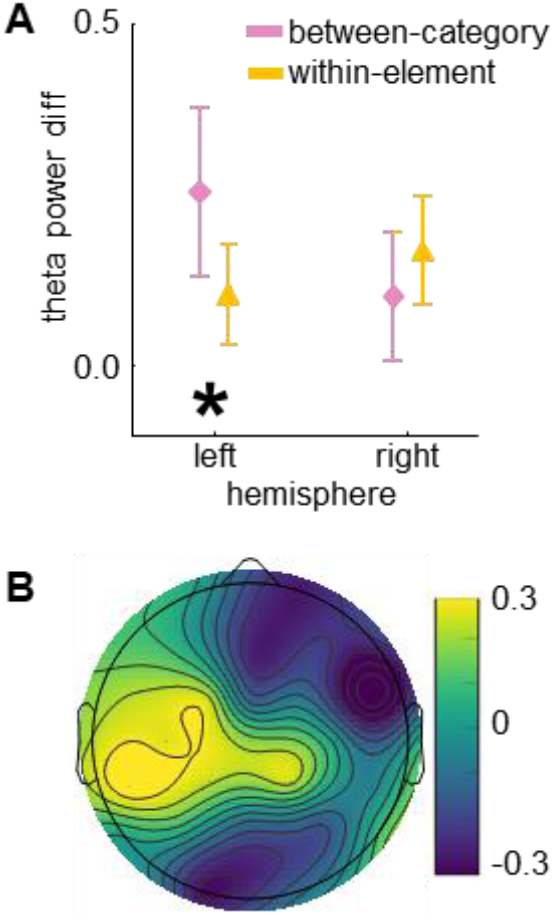
Comparison of the ‘within-element’ and ‘between-category’ theta memory effects. (A) The mean-error (mean ± SEM) plot of the power difference in the interval from 60 to 140 ms after the fixation onset. Significant result of the post-hoc test is indicated with an asterisk. (B) The power difference map: ‘between-category’ effect minus ‘within-element’ effect.

In sum, the correspondence of theta and eye movement results for between-category saccades and within-element saccades, which may be based on distinct mechanisms, indicates a diversity of relationships between theta activity, eye movements, and episodic memory formation, as will be discussed below.

## 4 Discussion

The present study set out to advance our current understanding of the relationship between eye movements and episodic memory encoding. In particular, we targeted the neural mechanisms that mediate the formation of coherent episodic memories during consecutive saccades to event elements during unrestricted viewing behavior. To this end, we applied a state-of-the-art analytical approach to EEG coregistered with eye movements during a free-viewing episodic memory task. Our approach provided a unique possibility to capture neural encoding mechanisms across eye movements at the level of gaze fixations while overcoming the confounding effects of sequential saccades on brain activity. We identified neural signatures that are associated with the buildup of episodic memories as a function of visual sampling behavior. The encoding of associations between event elements was accompanied by simultaneous modulation of eye movements and theta activity that were predictive of subsequent memory. This modulation likely reflects the binding of event elements into a coherent representation across eye movements. In addition, a subsequent memory effect was also observed for theta and alpha activity related to small scrutinizing saccades within elements. Although the association between exemplars within categories was incidental to the task, alpha activity related to between-exemplar saccades also proved predictive of later episodic remembering. Overall, we have identified the engagement of distinct neural mechanisms that may be the essential building blocks in forming episodic memory during unrestricted viewing.

Theta and alpha activity are considered to serve important but distinct functions of episodic memory formation: binding individual elements together into a whole (theta) and representing individual elements (alpha) (Hanslmayr et al., 2016). But it has not yet been established how these mechanisms are engaged when coherent episodic memories are formed in a piecemeal procedure across eye movements. In the current study, we aimed to fill this gap by tracking how these neural mechanisms subserve the buildup of episodic memories during unrestricted viewing behavior.

### 4.1 Saccade types that predict subsequent memory

It is well known that eye movement behavior during free viewing is important for the successful formation of episodic memory (for overviews, see Ryan et al., 2020; Wynn et al., 2019). We replicate and extend these findings by showing that the number of within-element saccades and between-category revisits positively correlate with performance in an associative memory task (Fig. 3C). Short within-element saccades scrutinize the visual features of single exemplars, increasing the visual sampling of different details, which leads to better subsequent memory of the stimuli (Liu et al., 2017; Loftus, 1972; Olsen et al., 2016).

As expected, the cumulative number of gaze transitions between categories during encoding increases memory performance. The between-category saccades presumably contribute to the binding of separate elements into a coherent representation of the event in which the hippocampus is involved (Herweg et al., 2020). Previous studies corroborate this idea by reporting that hippocampal activity and memory performance increase with the number of gaze fixations (Liu et al., 2017) and especially with the number of revisits (Kragel et al., 2021; Voss et al., 2011), as in our study.

In contrast to between-category saccades, the between-exemplar saccades did not predict subsequent memory performance. This was expected, as the memory test concerned between-category associations. This does, however, not mean that the between-exemplar saccades were not useful in forming a coherent event representation.

### 4.2 Theta and alpha modulations during the event encoding interval predict subsequent memory

First, the EEG analysis confirmed the presence of theta and alpha activity predictive of episodic remembering, extending previous work where the to-be-remembered material was presented centrally without the need for visual exploration via eye movements (Hanslmayr and Staudigl, 2014; Klimesch, 1996). Our results revealed that towards the end of the encoding interval (6-10 s), the theta power was higher for high than low subsequent event memory (Fig. 4A, C).

Consistent with previous research (Griffiths et al., 2019; Hanslmayr and Staudigl, 2014; Klimesch et al., 1996), we observed that high subsequent associative memory performance is related to a decrease in alpha power, mostly noticeable over the occipital areas (Fig. 4B). This decrease, however, became statistically significant only when we used a quadratic fit over time in the contrast analysis (Fig. 4D) while the subsequent memory effect for theta activity was linear.

Overall, in good agreement with the expected role of theta and alpha activity in episodic memory formation, we observe opposite effects predictive of subsequent memory in theta and alpha power. The theta increase and alpha decrease complement each other, reflecting the functioning of the hippocampal and cortical memory systems, respectively (Hanslmayr et al., 2016).

Additionally, we used the fBOSC method for oscillation detection (Seymour et al., 2022) to confirm the presence of true oscillatory activity in the data. In particular, the longest duration of oscillations (‘abundance’) in the encoding interval occurred around the alpha peak frequency. However, the abundance of alpha oscillatory activity did not predict subsequent memory performance (Fig. 4F). The duration of theta oscillations was shorter than the alpha oscillations; however, the theta oscillatory activity was predictive of memory performance. Oscillatory theta activity has been previously associated with memory performance (Hanslmayr et al., 2016), and our study replicates these previous findings. Alternatively, the duration of the theta oscillations may be associated with the eye movement rate since unrestricted eye movements are made at the theta frequency (Amit et al., 2017; Otero-Millan et al., 2008). Although oculomotor artifacts were eliminated from EEG during preprocessing, some residual artifacts at the rate of eye movements still could give rise to a pattern that appears as theta oscillations. However, the observed eye movement rate was much higher in low than high memory performance. Consequently, this eliminates the possibility that the eye movement rhythmicity could mechanistically determine the long duration of theta oscillatory activity, as well as the large power of theta oscillations that characterized high memory performance. Instead, we propose that the ‘long’ duration of memory-related theta oscillations is associated with the modulation of ongoing or saccade-induced theta oscillations (see below). Such oscillations may overlap, which would require correction by deconvolution. Perhaps this can be done by combining the oscillation detection methods (such as those used in 2.5.4), providing latencies of oscillation onsets, with the deconvolution technique. However, solutions to this intricate issue await method development far beyond the scope of the present study.

### 4.3 Fixation-related EEG predicts subsequent memory

The fixation-related EEG corroborates the findings reported at the event level. We observed reliably higher theta and lower alpha power for high than low memory performance (Fig. 5). In line with the findings at the event level, theta and alpha memory effects became greater over time of the encoding interval (Fig. 5B, F), which may reflect the buildup of an event representation across a sequence of eye movements. The predominantly anterior topography of the theta memory effect (Fig. 5D) suggests the contribution of frontal midline theta oscillations, which are related to intrinsic episodic memory processes (Hsieh and Ranganath, 2014). The alpha memory effect was predominantly distributed over the left hemisphere.

Memory processes in natural viewing are much more complex and evolve at a much higher rate than is considered by the design of traditional paradigms with fixed stimulus presentation. Depending on the fixation location, learning with unrestricted eye movements may involve encoding *and* retrieval, just as later remembering may trigger retrieval *and* encoding processes (Kragel and Voss, 2022). These dynamic memory processes across eye movements produce multiple effects on the underlying brain activity. The ongoing memory-related theta and alpha oscillations described above interfere with the activity elicited by saccade execution and information acquisition at new fixation locations. Each saccade evokes a broadband neural response (Dimigen et al., 2009; Ossandón et al., 2010) that lasts longer than 200-300 ms typical for fixation duration in natural viewing and therefore overlaps each other (Dimigen and Ehinger, 2021; Dimigen et al., 2011; Nikolaev et al., 2016). Moreover, each saccade aligns the phase of ongoing oscillations observed in the visual cortex of monkeys (Ito et al., 2011; Rajkai et al., 2008), in the human and macaque hippocampus (Hoffman et al., 2013), as well as in the scalp EEG in humans (Nikolaev et al., 2016). Such phase resetting supports successful memory formation (Jutras et al., 2013; Kota et al., 2020; Kragel et al., 2020) by synchronizing brain areas and optimizing the processing and encoding of visual information (Jutras and Buffalo, 2010; Voss et al., 2017). Thus, the neural activity during unrestricted viewing behavior constitutes a combination of overlapping evoked responses and the phase-reset of ongoing oscillations (Ito et al., 2011). In such a situation, mapping external stimuli, mental processes, and neural activity is not straightforward, as in the traditional stimulus-response paradigm. In particular, ongoing oscillations may simultaneously represent the current and previous memory items (Herweg et al., 2020), and overlapping responses evoked by a series of eye movements may represent information about several previous fixation locations. The relative contribution of saccade-evoked responses and phase-reset oscillations to the resulting neural activity at fixation is challenging to estimate. Accordingly, the question remains open as to how the ongoing memory-related oscillations interact with the fixation-related activity representing visual processing at the current fixation location. It is clear, however, that neural activity time-locked to fixation onsets in a free-viewing memory task may reflect not only information acquisition at the currently fixated location but also preceding mental representations and more sustained higher-order memory processes.

The time window in the fixation-related EEG analysis was centered on the first power peak about 100 ms after the fixation onset. The latency of this peak coincides with the latency of lambda activity, which is associated with early perceptual processes at gaze fixation (Kazai and Yagi, 2003; Ossandón et al., 2010; Ries et al., 2016; Thickbroom et al., 1991). However, given that observed memory-related EEG effects have topographies that do not overlap with the typical occipital prominence of lambda activity (Ossandón et al., 2010; Thickbroom et al., 1991), it seems unlikely that the perceptual lambda activity would explain them. This interpretation is corroborated by previous research, where a comparable memory-related peak latency has been reported for frontal theta power (Adam et al., 2018). Ongoing theta oscillations supposedly drive perceptual sampling by organizing sequential inputs: they represent several items within the same cycle, where the current item is strongly represented and previous items are concurrently represented in a weaker form (Herweg et al., 2020). Such theta oscillations may thus be further modulated by fixation-related activity, where not only the currently fixated information is represented but also previously visited elements from the ongoing event memorization process (as well as previous events). Thus, our time window may capture individual perceptual moments at fixations and the sum of active and latent representations accumulated over a series of eye movements.

### 4.4 EEG memory effects after saccades between event elements

Fascinating results were obtained when analyzing EEG in fixation intervals after different saccade types during the 10-s encoding interval: fixations following between-category, between-exemplar, and within-element saccades. In our paradigm, between-category saccades supported associations between event elements that were directly targeted in the episodic memory task. The theta power was higher over the left hemisphere for high than low subsequent memory after between-category revisits (Fig. 6D, E), and the cumulative number of such revisits predicted subsequent memory (Fig. 3C). The alpha power was lower for high than low memory at fixations after between-exemplar saccades, i.e., the associations that were not tested (Fig. 7). The cumulative number of such gaze transitions did not predict subsequent memory (Fig. 3C).

It is conceivable that gaze transitions between categories link the event elements together, creating a holistic representation of the entire event. These gaze transitions are thus critical to episodic memory as they support the binding of information experienced separately in space and time (Cohen and Eichenbaum, 1993; Voss et al., 2017). Previous research has shown that this binding mechanism relies on the transfer of information between distant brain regions and is promoted by synchronizing their activity at a theta rhythm (Clouter et al., 2017; Fell and Axmacher, 2011). Our results fit well with this previous literature. We observed high theta power after between-category saccades, predictive of high event memory, suggesting the involvement of theta in the binding of elements into a coherent episodic representation. It may be the case that theta reflects an optimized oculomotor control that guides this binding mechanism. In fact, it has been shown that the theta rhythm also underlies viewing behavior optimal for memory formation (Herweg et al., 2020; Jutras and Buffalo, 2010; Voss et al., 2017).

The crucial role of the hippocampal theta oscillations in memory-related gaze behavior has recently been demonstrated using intracranial EEG recording combined with eye tracking in free-viewing memory tasks (Kragel et al., 2021; Kragel et al., 2020). Kragel and colleagues (2021) described how revisits support the efficiency of visual sampling in the formation of memory for scenes. Hippocampal theta power decreased before revisits and increased after revisits, indicating rapid switches between memory-based saccade guidance and perceptual processing at fixation. Thus, the theta increase after revisits may reflect the mechanism binding information encoded at multiple fixations into a coherent representation. The present study supports the crucial role of revisits in successful memory formation: the number of between-category revisits and succeeding theta power predicts subsequent associative memory performance. These results demonstrate the feasibility of applying our approach in free-viewing studies of episodic memory in broad cohorts of healthy participants.

As expected, the cumulative number of between-exemplar saccades that were incidental to the task did not predict subsequent memory. Nevertheless, alpha power at fixations after these saccades was lower for high than low subsequent memory (Fig. 7). Although this alpha effect thus reflects a mechanism that supports successful memory formation, its exact nature remains unclear. For example, in episodic memory encoding, alpha activity indicates cortical information processing during the perception of an event (Jensen and Mazaheri, 2010; Klimesch, 2012), and its decrease predicts the successful formation of episodic memory (Hanslmayr et al., 2016). As discussed above, alpha desynchronization is thought to increase the capacity of local cell ensembles to process information by reducing local synchronization (Hanslmayr et al., 2016). Thus, the occipital predominance of memory-related alpha decrease in our visual memory task may reflect an increase in the capacity of cell ensembles in the visual system, which is beneficial for category-specific processing. Visual perception in our task involves discriminating between two similar exemplars from the same category, and the between-exemplar saccades may facilitate a comparison of the two exemplars. Alpha activity has been shown to predict behavioral performance in visual discrimination tasks (Hanslmayr et al., 2005; Nelli et al., 2017). Thus, the here observed alpha decrease after between-exemplar saccades may indicate efficient discrimination of two similar exemplars that promote successful encoding.

### 4.5 EEG memory effects after saccades within event elements

Within-element saccades are the only saccade type for which memory-related effects were found in both theta and alpha bands. They had the same direction in both frequency bands: high power predicted high memory. After within-element saccades, alpha power was higher for high than low subsequent memory performance over the right hemisphere (Fig. 8E-G), whereas theta power was higher for high than low memory performance over the frontal and central areas. Short within-element saccades that serve to scrutinize the visual features of images are known to increase visual sampling, which leads to better performance on subsequent item recognition tests (Loftus, 1972; Olsen et al., 2016). Accordingly, it is conceivable that a higher number of within-element saccades helps establish strong individual element representations to form part of multi-element event representations. Increased visual sampling results in a high memory load, reflected in high frontal theta power (Jensen and Tesche, 2002; Sederberg et al., 2006). In our paradigm, an increased memory load across encoding time corresponds to the accumulation of more visual details about event exemplars, which is favorable for later recall. Alternatively, frontal theta power has also been associated with detecting interference during selective retrieval in episodic memory (Ferreira et al., 2014; Kerrén et al., 2021; Staudigl et al., 2010) and during multiple-list learning in working memory (Kliegl and Bauml, 2021). In the present study, encoding exemplars from multiple events may be accompanied by interference from exemplars of the same categories in previously acquired events. Therefore, detecting interference may be crucial for encoding distinct event memories and may trigger adaptive inhibition of exemplars from previous events, ensuring that each event is stored separately to promote task performance. It is also conceivable that the frontal theta reflects the complementary effects of interference detection and memory load, as high memory load can lead to increased interference. The topography of the theta memory effect for within-element saccades differs from that of the between-category saccades, which was more prominent over the left hemisphere (Fig. 9), suggesting that those two theta effects represent distinct neural mechanisms.

There is overwhelming evidence that task-related increase in alpha power often corresponds to inhibitory neural processes (Jensen and Mazaheri, 2010). In studies of competitive memory retrieval, it has been demonstrated that high alpha power is associated with the inhibition of goal-irrelevant memories (Fellner et al., 2020; Waldhauser et al., 2012). Considering the likely involvement of both encoding and retrieval processes during free-viewing memory formation (Kragel and Voss, 2022), the observed alpha power increase may indicate inhibition of competing information (within and across events) that may interfere with building a coherent event representation across momentary sequences of visual sampling.

## 5 Conclusions

We used a novel method of EEG-eye movement coregistration to examine episodic memory encoding in unrestricted viewing with a precision unavailable thus far. Our findings confirm eye movements’ crucial role in forming episodic memory (Voss et al., 2017). We demonstrate that the functional specialization of eye movements manifested in various types of saccades is associated with different perceptual and cognitive processes, as evidenced by fixation-related theta and alpha EEG power. Episodic memory about an event is constructed through rapid switching between these processes in a sequence of fixations. These processes may indicate the complementary functioning of the hippocampal theta system for binding episodes and the cortical alpha system for representing the content of these episodes (Hanslmayr et al., 2016). Our study extends previous knowledge by elucidating how these systems engage across eye movements in the service of building coherent episodic memories for personal experiences.

## Supporting information

Supplementary material_Revis

## Acknowledgments

We thank Benedikt Ehinger for his help with deconvolution and Axel Ekström for his assistance with data collection. We are grateful to Marcus Nyström and Diederick Niehorster for their help in developing the EEG-eye movement coregistration system.

## Data and code availability statement

The data and main codes used in this article are available at Open Science Framework: https://osf.io/nad4y/?view_only=df274a4eaecc4ef6b9ec5053a0d2b1f0

## CRediT authorship contribution statement

**Andrey R. Nikolaev**: Conceptualization, Methodology, Software, Formal analysis, Writing - Original Draft, Writing - Review & Editing. **Inês Bramão**: Conceptualization, Methodology, Writing - Review & Editing. **Roger Johansson**: Conceptualization, Methodology, Writing - Review & Editing. **Mikael Johansson**: Conceptualization, Methodology, Writing - Review & Editing, Supervision, Funding acquisition.

