## Supplementary material_Revis for "Episodic memory formation in unrestricted viewing"

#### Events in which fixations were detected on only two of the three categories

We investigated the characteristics of events in which fixations were detected on only two of the three categories. Across the 20 participants who had such events, the mean memory score was about the same for the event with two (35.9, SD = 12.7) and three (36.5, SD = 8.4) fixated categories ($t(19) = 0.3$, $p = .78$). The distribution of such events across the six blocks was not uniform (Chi-Square = 9.6, df = 3, $p = .02$), and the number of such events was smallest in the first block and largest in the fifth block (Fig. S1). Notably, the duration of 'bad eye movement intervals' (i.e., when the eye-tracking signal was lost) was much longer for such events (1950, SD = 1215 ms) compared to events with three (180, SD = 68 ms) fixated categories ($t(18) = 6.5$, $p < .001$). These findings indicate that events with only two fixated categories primarily occurred due to losses of the eye-tracking signal, which mainly developed towards the end of the experiment (likely due to participants moving away or closing their eye lids due to fatigue or reduced vigilance.

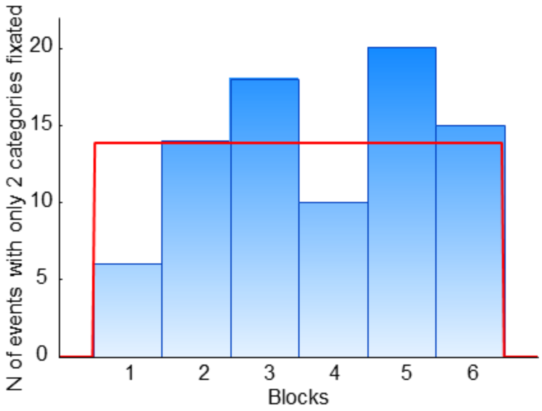


Fig. S1. The distribution of events in which fixations were detected on only two of the three categories across six blocks of the experiment. The red line indicates the fit with the expected uniform distribution.

### Memory score regardless of confidence

We evaluated the contribution of confidence to memory scores used in the main analyses. To that end, we applied a repeated measures ANOVA on the memory scores that were not weighted by confidence with the factors of Event (nine levels) and Block (six levels) (Fig. S2). There were significant main effects of Event ($F(8, 216) = 2.9$, $p = .01$, $\varepsilon = .68$) and Block ($F(5, 135) = 9.9$, $p < .001$, $\varepsilon = .95$)), but no interaction between them. To characterize the effects of Event and Block, we used planned comparisons with the linear and quadratic contrasts. The contrast analysis revealed significant effects for the linear contrast for Block ($F(1, 27) = 34.7$, $p < .001$) and for the quadratic contrast for Event ($F(1, 27) = 9.3$, $p = .005$), but not for the linear contrast for Event ($F(1, 27) = 0.5$, $p = .5$). These results were qualitatively the same as those for scores weighted by confidence. They indicate a prominent learning effect during the experiment, as well as possible primacy and recency effects in time course of a block.

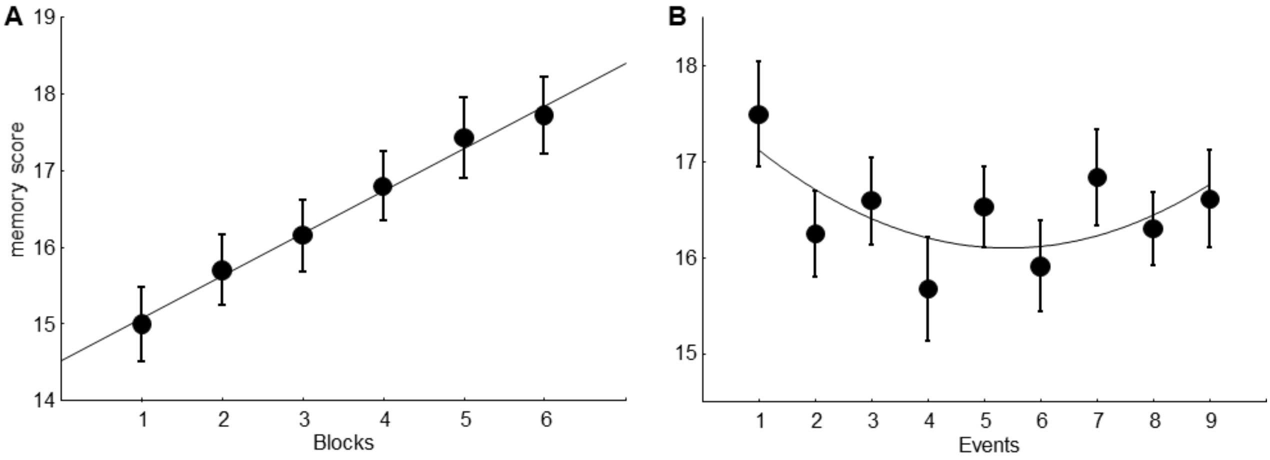


Fig. S2. Memory performance (regardless of confidence) in the time course of an experiment (A) and a block (B). The thin lines indicate significant linear (A) and quadratic (B) fits.

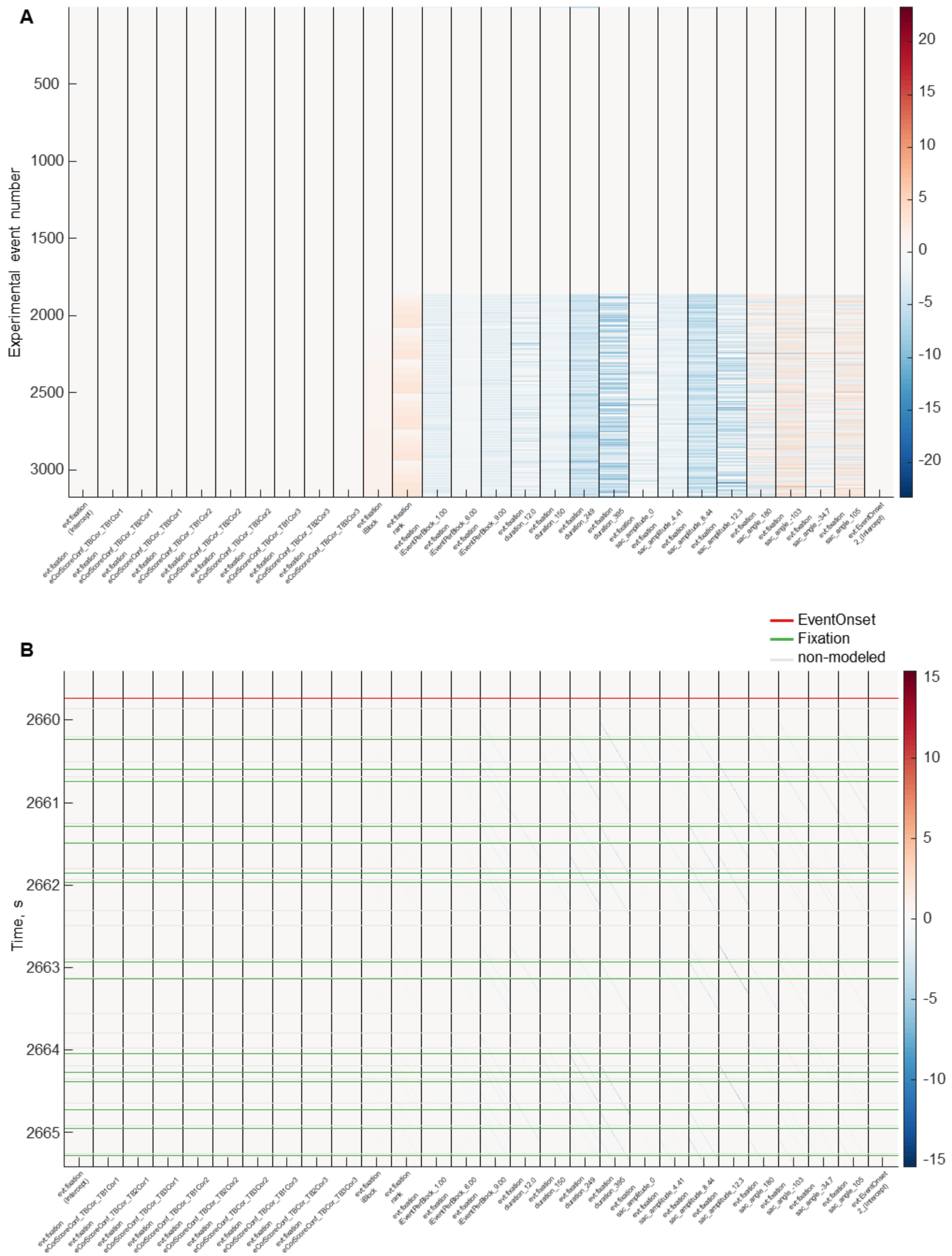


Fig. S3. (A) The design matrix for one participant. The entire design matrix is shown for the formula, which included 3 Memory bins, 3 Rank bins, and all eye movement and ordering predictors (the results of this analysis are described in 3.4.1). The large number of experimental events are defined by the number of fixations. (B) The time-expanded design matrix: it is very sparse, so only a 5-s fragment is shown. The onsets of events, fixations and non-modelled intervals are shown by horizontal lines. The design matrices were obtained with the function *uf_plotDesignmat* of the Unfold toolbox.

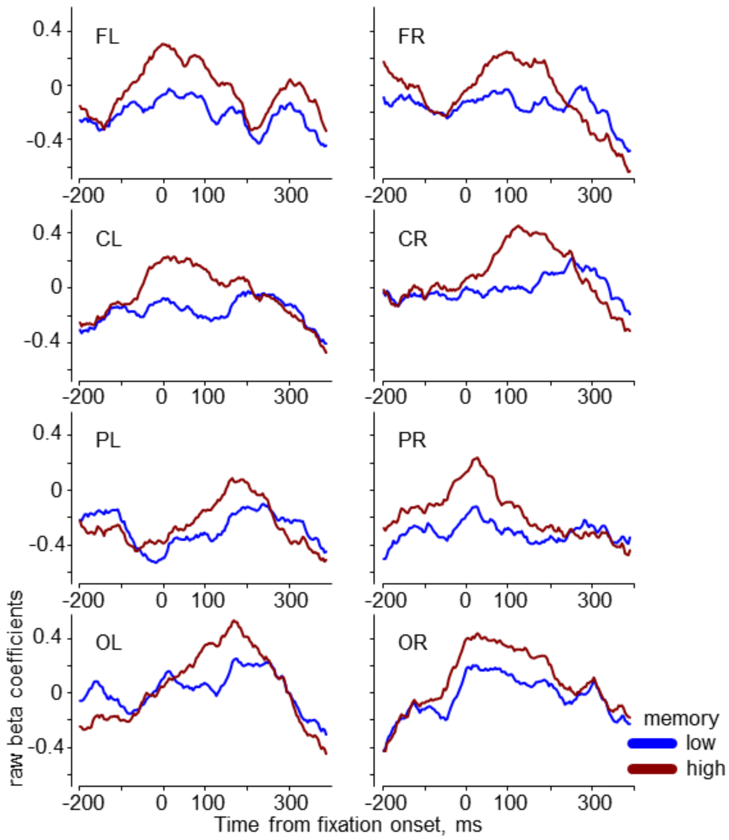


Fig. S4. The raw beta coefficients after deconvolution, but before addition of the intercept and the marginals of all continuous and spline predictors (i.e., before applying the functions *uf_predictContinuous* and *uf_addmarginal* of the Unfold toolbox). The plot presents the grand-averaged (N=28) theta power in the low and high memory conditions for rank bin 3 (see Section 3.4.1), i.e., it is analogous to Fig. 5A, to which the intercept and the marginals have been added. Note that the waveforms oscillate around zero, but the difference between high and low memory is preserved.

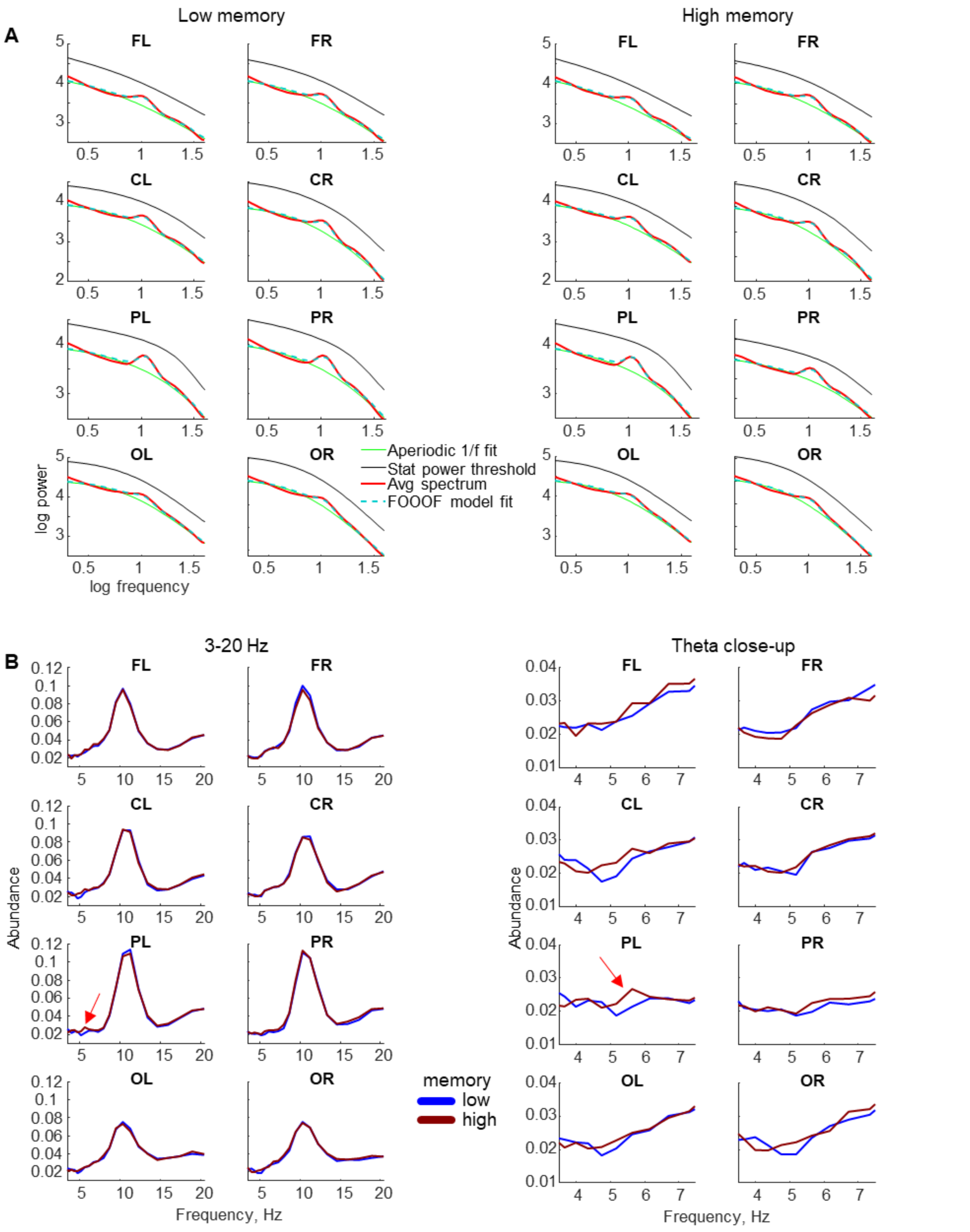


Fig. S5. (A) Log-log plot of the EEG power spectrum averaged across 28 participants for low (left) and high (right) subsequent memory performance for 8 ROIs (frontal, central, parietal, occipital for the left and right hemisphere: FL, FR, CL, CR, PL, PR, OL, OR). (B) The duration of the rhythmic episode relative to the length of the analyzed segment ('abundance') averaged across 28 participants for 8 ROIs for low and high memory performance. (left) Abundance for the frequency range 3-20 Hz. (right) Abundance in the theta band (close-up of the left plot). The arrows indicate the maximal abundance increase for high memory performance.

### Evaluation of deconvolution

We evaluated the general effects of deconvolution correction on the power waveforms. Specifically, we took as a reference the condition with a prominent theta memory effect for rank bin 3 after deconvolution and after the addition of intercept and covariates, i.e., after complete deconvolution correction of the overlapping effects. The grand-averaged (N=28) power waveforms for this condition are shown in Fig. 5A. We compared the shapes of these waveforms with the same waveforms, but without correction of overlapping effects as well as eye movement and ordering covariates (Fig. S6A). To achieve this, we included in the formula only the categorical predictor of memory performance, but no eye movement or ordering predictors. Then, instead of deconvolution, we used a massive univariate linear model (the function *uf_glmfit_nodc* of the Unfold toolbox). The intercept was added to memory predictors.

To explore the effect of deconvolution correction we compared Fig. 5A and Fig. S6A. At a first glance the plots may look similar, and the memory effect (higher power for high than low memory) is noticeable in both plots. However, in Fig. S6A (without deconvolution) the power values are more negative. This is highlighted in in Fig. S6B and C, where we plotted the power waveforms with and without deconvolution for the two ROIs: FL and OR. This large difference in power values may result from correcting the effect of fixation rank, which is known to produce a systematic effect on the fixation-related EEG during time course of a free-viewing trial with multiple eye movements (Kamienkowski et al., 2018; Nikolaev et al. 2022). This effect appears to have been eliminated by including fixation rank to the deconvolution model. Another effect of deconvolution correction is visible at both edges of the power waveforms, where the overlaps with the previous and following eye movements should be maximal. After deconvolution, these edges become steeper, suggesting that removing the overlap makes the oscillatory activity more distinct.

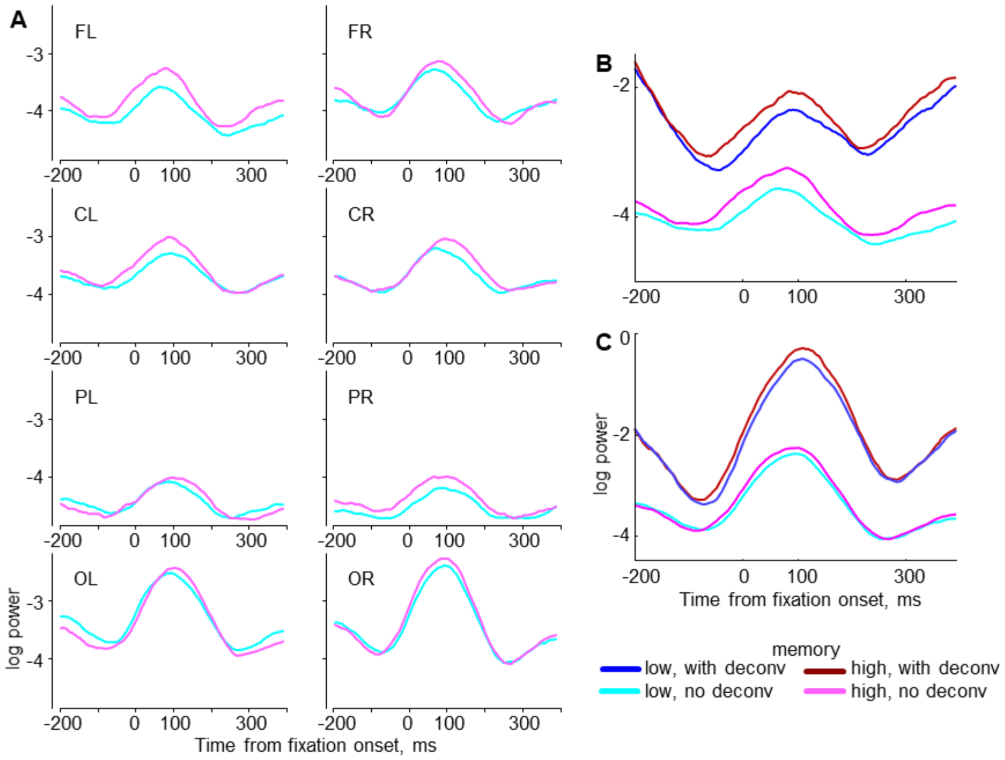


Fig. S6. The effect of deconvolution correction. (A) The grand-averaged (N=28) theta power in the low and high memory conditions for rank bin 3 *without* deconvolution correction. This plot is analogous to the plot in Fig. 5A with deconvolution correction. (B) The power waveforms with and without deconvolution for the left frontal (FL) ROI. (C) The power waveforms with and without deconvolution for the right occipital (OR) ROI.

### Table S1

The number of fixations used in the deconvolution modeling of each memory and rank bin condition per participant: mean (range, SD) across 28 participants.

| | low memory | medium memory | high memory |
|---|---|---|---|
| Fixation rank bin 1 | 128.7 (77-217, 34.0) | 113.8 (65-207, 32.9) | 103.4 (50-148, 25.8) |
| Fixation rank bin 2 | 116.0 (71-200, 32.2) | 103.1 (54-194, 32.5) | 93.2 (43-136, 25.0) |
| Fixation rank bin 3 | 121.7 (75-208, 32.6) | 108.6 (60-202, 33.6) | 98.2 (48-142, 25.6) |

### Table S2

The complete ANOVA results of all EEG analyses. The corresponding sections of Results are indicated before each sub-table.

**Factors**
MEM: Memory (Low vs. High)
ROI: Regions of interest (Frontal, Central, Parietal, Occipital)
HEMI: Hemisphere (Left vs. Right)
TIME_BIN: Time (5 bins)
RANK_BIN: Fixation Rank (3 bins)
GAZE: Gaze transition status (First visits vs. Revisit)

**Statistical abbreviations**
df: degrees of freedom;
F: F-value; p: p-level;
H-F – Epsilon: Huynh-Feldt epsilon;
H-F - Adj. p: Huynh-Feldt adjusted p-level.
Significant (p < .05) differences are marked in red.

Section 3.3.1: The theta and alpha power at the event level in 5 time bins

| Theta | df | F | p | H-F - Epsilon | H-F - Adj. p |
|---|---|---|---|---|---|
| MEM | 1, 27 | 1.12459 | 0.298326 | 1 | 0.298326 |
| ROI | 3, 81 | 12.71871 | 0.000001 | 0.628526 | 0.000046 |
| HEMI | 1, 27 | 5.10415 | 0.03215 | 1 | 0.03215 |
| TIME_BIN | 4, 108 | 7.10751 | 0.000041 | 0.752346 | 0.000263 |
| MEM*ROI | 3, 81 | 0.37839 | 0.768827 | 0.939497 | 0.756473 |
| MEM*HEMI | 1, 27 | 0.87519 | 0.357814 | 1 | 0.357814 |
| ROI*HEMI | 3, 81 | 0.60272 | 0.615099 | 0.760981 | 0.571737 |
| MEM*TIME_BIN | 4, 108 | 4.8759 | 0.001184 | 0.986292 | 0.001258 |
| ROI*TIME_BIN | 12, 324 | 2.03016 | 0.021328 | 0.687894 | 0.042159 |
| HEMI*TIME_BIN | 4, 108 | 1.66279 | 0.163936 | 1 | 0.163936 |
| MEM*ROI*HEMI | 3, 81 | 0.17699 | 0.911695 | 0.70325 | 0.849096 |
| MEM*ROI*TIME_BIN | 12, 324 | 0.8881 | 0.559429 | 0.870925 | 0.548306 |
| MEM*HEMI*TIME_BIN | 4, 108 | 1.0219 | 0.399412 | 1 | 0.399412 |

| | | | | | |
|---|---|---|---|---|---|
| ROI*HEMI*TIME_BIN | 12, 324 | 0.89961 | 0.547667 | 0.653866 | 0.516218 |
| MEM*ROI*HEMI*TIME_BIN | 12, 324 | 1.18714 | 0.29092 | 0.711069 | 0.305696 |
| **Alpha** | **df** | **F** | **p** | **H-F - Epsilon** | **H-F - Adj. p** |
| MEM | 1, 27 | 0.58551 | 0.450798 | 1.000000 | 0.450798 |
| ROI | 3, 81 | 14.76875 | 0.000000 | 0.624375 | 0.000013 |
| HEMI | 1, 27 | 5.94842 | 0.021585 | 1.000000 | 0.021585 |
| TIME_BIN | 4, 108 | 11.70625 | 0.000000 | 0.746661 | 0.000002 |
| MEM*ROI | 3, 81 | 1.03703 | 0.380729 | 0.810906 | 0.371357 |
| MEM*HEMI | 1, 27 | 0.15812 | 0.694020 | 1.000000 | 0.694020 |
| ROI*HEMI | 3, 81 | 10.46048 | 0.000007 | 0.783026 | 0.000050 |
| MEM*TIME_BIN | 4, 108 | 0.81603 | 0.517640 | 0.875587 | 0.504272 |
| ROI*TIME_BIN | 12, 324 | 6.34637 | 0.000000 | 0.610304 | 0.000001 |
| HEMI*TIME_BIN | 4, 108 | 1.85095 | 0.124383 | 0.768907 | 0.142899 |
| MEM*ROI*HEMI | 3, 81 | 0.85977 | 0.465495 | 0.709187 | 0.434648 |
| MEM*ROI*TIME_BIN | 12, 324 | 0.99100 | 0.457148 | 0.750434 | 0.448018 |
| MEM*HEMI*TIME_BIN | 4, 108 | 1.89004 | 0.117383 | 0.671416 | 0.144860 |
| ROI*HEMI*TIME_BIN | 12, 324 | 1.04643 | 0.405734 | 0.544835 | 0.399018 |
| MEM*ROI*HEMI*TIME_BIN | 12, 324 | 0.95936 | 0.487793 | 0.642961 | 0.467074 |

Section 3.3.2: Abundance in the theta and alpha bands

| **Theta** | **df** | **F** | **p** | **H-F - Epsilon** | **H-F - Adj. p** |
|---|---|---|---|---|---|
| MEM | 1, 27 | 4.752328 | 0.038151 | 1.000000 | 0.038151 |
| ROI | 3, 81 | 9.226176 | 0.000026 | 0.912619 | 0.000051 |
| HEMI | 1, 27 | 1.570906 | 0.220821 | 1.000000 | 0.220821 |
| MEM*ROI | 3, 81 | 0.822940 | 0.484975 | 0.855769 | 0.469560 |
| MEM*HEMI | 1, 27 | 2.134082 | 0.155597 | 1.000000 | 0.155597 |
| ROI*HEMI | 3, 81 | 1.686885 | 0.176330 | 1.000000 | 0.176330 |
| MEM*ROI*HEMI | 3, 18 | 2.285503 | 0.084979 | 0.983363 | 0.086143 |
| **Alpha** | **df** | **F** | **p** | **H-F - Epsilon** | **H-F - Adj. p** |
| MEM | 1, 27 | 0.35153 | 0.558183 | 1.000000 | 0.558183 |
| ROI | 3, 81 | 17.97291 | 0.000000 | 0.924116 | 0.000000 |
| HEMI | 1, 27 | 0.10932 | 0.743469 | 1.000000 | 0.743469 |
| MEM*ROI | 3, 81 | 0.90369 | 0.443123 | 0.914416 | 0.436257 |
| MEM*HEMI | 1, 27 | 0.27432 | 0.604719 | 1.000000 | 0.604719 |
| ROI*HEMI | 3, 81 | 0.54331 | 0.654042 | 0.763580 | 0.607294 |
| MEM*ROI*HEMI | 3, 18 | 1.38044 | 0.254676 | 0.848102 | 0.257960 |

Section 3.4.1: The theta and alpha power at the fixation level in 3 fixation rank bins

| **Theta** | **df** | **F** | **p** | **H-F - Epsilon** | **H-F - Adj. p** |
|---|---|---|---|---|---|
| MEM | 1, 27 | 1.98104 | 0.170690 | 1.000000 | 0.170690 |
| RANK_BIN | 2, 54 | 1.43214 | 0.247721 | 0.873103 | 0.248514 |
| ROI | 3, 81 | 37.64292 | 0.000000 | 0.863174 | 0.000000 |
| HEMI | 1, 27 | 5.69137 | 0.024327 | 1.000000 | 0.024327 |

| | | | | | |
|---|---|---|---|---|---|
| MEM*RANK_BIN | 2, 54 | 3.54810 | 0.035665 | 0.955972 | 0.037888 |
| MEM*ROI | 3, 81 | 3.59764 | 0.016991 | 0.856194 | 0.022923 |
| RANK_BIN*ROI | 6, 162 | 3.88867 | 0.001180 | 0.790302 | 0.003076 |
| MEM*HEMI | 1, 27 | 2.18284 | 0.151128 | 1.000000 | 0.151128 |
| RANK_BIN*HEMI | 2, 54 | 2.27113 | 0.112968 | 0.690177 | 0.132143 |
| ROI*HEMI | 3, 81 | 6.30256 | 0.000675 | 0.829177 | 0.001556 |
| MEM*RANK_BIN*ROI | 6, 162 | 0.33346 | 0.918508 | 0.902759 | 0.904078 |
| MEM*RANK_BIN*HEMI | 2, 54 | 1.57910 | 0.215532 | 0.950429 | 0.216736 |
| MEM*ROI*HEMI | 3, 81 | 1.83695 | 0.147001 | 0.726759 | 0.165095 |
| RANK_BIN*ROI*HEMI | 6, 162 | 1.89681 | 0.084341 | 0.518206 | 0.134284 |
| MEM*RANK_BIN*ROI*HEMI | 6, 162 | 0.45896 | 0.837816 | 0.704742 | 0.775901 |
| **Alpha** | **df** | **F** | **p** | **H-F - Epsilon** | **H-F - Adj. p** |
| MEM | 1, 27 | 1.20868 | 0.281300 | 1.000000 | 0.281300 |
| RANK_BIN | 2, 54 | 0.06700 | 0.935271 | 0.927807 | 0.924283 |
| ROI | 3, 81 | 25.39978 | 0.000000 | 0.814811 | 0.000000 |
| HEMI | 1, 27 | 15.08479 | 0.000601 | 1.000000 | 0.000601 |
| MEM*RANK_BIN | 2, 54 | 0.34925 | 0.706798 | 0.918491 | 0.688847 |
| MEM*ROI | 3, 81 | 0.93205 | 0.429158 | 0.853179 | 0.418074 |
| RANK_BIN*ROI | 6, 162 | 2.10381 | 0.055526 | 0.632735 | 0.089155 |
| MEM*HEMI | 1, 27 | 3.35410 | 0.078094 | 1.000000 | 0.078094 |
| RANK_BIN*HEMI | 2, 54 | 0.20542 | 0.814938 | 0.850539 | 0.779640 |
| ROI*HEMI | 3, 81 | 2.67114 | 0.052921 | 0.830843 | 0.064551 |
| MEM*RANK_BIN*ROI | 6, 162 | 1.10365 | 0.362479 | 0.915219 | 0.362122 |
| MEM*RANK_BIN*HEMI | 2, 54 | 4.99166 | 0.010252 | 0.981776 | 0.010707 |
| MEM*ROI*HEMI | 3, 81 | 0.16842 | 0.917400 | 0.590664 | 0.820070 |
| RANK_BIN*ROI*HEMI | 6, 162 | 0.17737 | 0.982662 | 0.592042 | 0.935389 |
| MEM*RANK_BIN*ROI*HEMI | 6, 162 | 2.37113 | 0.031917 | 0.837847 | 0.042179 |

Section 3.4.2.1: The theta and alpha power after between-category saccades

| Theta | df | F | p | H-F - Epsilon | H-F - Adj. p |
|---|---|---|---|---|---|
| GAZE | 1, 27 | 4.99003 | 0.033973 | 1.000000 | 0.033973 |
| MEM | 1, 27 | 3.07101 | 0.091051 | 1.000000 | 0.091051 |
| ROI | 3, 81 | 36.86924 | 0.000000 | 0.812750 | 0.000000 |
| HEMI | 1, 27 | 1.88755 | 0.180778 | 1.000000 | 0.180778 |
| GAZE*MEM | 1, 27 | 0.80121 | 0.378641 | 1.000000 | 0.378641 |
| GAZE*ROI | 3, 81 | 4.17547 | 0.008414 | 0.943762 | 0.009774 |
| MEM*ROI | 3, 81 | 0.45805 | 0.712352 | 0.803668 | 0.670688 |
| GAZE*HEMI | 1, 27 | 1.23244 | 0.276723 | 1.000000 | 0.276723 |
| MEM*HEMI | 1, 27 | 0.11311 | 0.739235 | 1.000000 | 0.739235 |
| ROI*HEMI | 3, 81 | 4.72281 | 0.004351 | 0.843829 | 0.007214 |
| GAZE*MEM*ROI | 3, 81 | 0.17072 | 0.915875 | 0.689483 | 0.850305 |
| GAZE*MEM*HEMI | 1, 27 | 11.75056 | 0.001965 | 1.000000 | 0.001965 |
| GAZE*ROI*HEMI | 3, 81 | 0.21054 | 0.888832 | 0.673849 | 0.813081 |
| MEM*ROI*HEMI | 3, 81 | 0.22384 | 0.879567 | 0.763393 | 0.828441 |

| GAZE*MEM*ROI*HEMI | 3, 81 | 0.40585 | 0.749191 | 0.916273 | 0.732005 |
|---|---|---|---|---|---|
| **Alpha** | **df** | **F** | **p** | **H-F - Epsilon** | **H-F - Adj. p** |
| GAZE | 1, 27 | 1.71127 | 0.201848 | 1.000000 | 0.201848 |
| MEM | 1, 27 | 3.08600 | 0.090306 | 1.000000 | 0.090306 |
| ROI | 3, 81 | 27.20665 | 0.000000 | 0.817113 | 0.000000 |
| HEMI | 1, 27 | 10.72964 | 0.002894 | 1.000000 | 0.002894 |
| GAZE*MEM | 1, 27 | 0.46155 | 0.502685 | 1.000000 | 0.502685 |
| GAZE*ROI | 3, 81 | 1.00819 | 0.393533 | 0.933297 | 0.390103 |
| MEM*ROI | 3, 81 | 0.48870 | 0.691094 | 0.900391 | 0.671676 |
| GAZE*HEMI | 1, 27 | 0.00913 | 0.924584 | 1.000000 | 0.924584 |
| MEM*HEMI | 1, 27 | 0.29870 | 0.589186 | 1.000000 | 0.589186 |
| ROI*HEMI | 3, 81 | 2.62824 | 0.055786 | 0.758030 | 0.073410 |
| GAZE*MEM*ROI | 3, 81 | 2.12673 | 0.103226 | 0.925566 | 0.108567 |
| GAZE*MEM*HEMI | 1, 27 | 0.72489 | 0.402032 | 1.000000 | 0.402032 |
| GAZE*ROI*HEMI | 3, 81 | 2.67758 | 0.052504 | 0.695817 | 0.075215 |
| MEM*ROI*HEMI | 3, 81 | 0.42031 | 0.738916 | 0.764796 | 0.686231 |
| GAZE*MEM*ROI*HEMI | 3, 81 | 0.21561 | 0.885311 | 0.573545 | 0.773848 |

Section 3.4.2.2: The theta and alpha power after between-exemplar saccades

| **Theta** | **df** | **F** | **p** | **H-F - Epsilon** | **H-F - Adj. p** |
|---|---|---|---|---|---|
| GAZE | 1, 27 | 2.09724 | 0.159079 | 1.000000 | 0.159079 |
| MEM | 1, 27 | 0.30676 | 0.584231 | 1.000000 | 0.584231 |
| ROI | 3, 81 | 37.35610 | 0.000000 | 0.854102 | 0.000000 |
| HEMI | 1, 27 | 7.53878 | 0.010612 | 1.000000 | 0.010612 |
| GAZE*MEM | 1, 27 | 1.46028 | 0.237364 | 1.000000 | 0.237364 |
| GAZE*ROI | 3, 81 | 1.85834 | 0.143227 | 0.892883 | 0.150303 |
| MEM*ROI | 3, 81 | 1.88135 | 0.139274 | 0.892869 | 0.146452 |
| GAZE*HEMI | 1, 27 | 0.25784 | 0.615731 | 1.000000 | 0.615731 |
| MEM*HEMI | 1, 27 | 3.23868 | 0.083104 | 1.000000 | 0.083104 |
| ROI*HEMI | 3, 81 | 6.80139 | 0.000380 | 0.816460 | 0.001025 |
| GAZE*MEM*ROI | 3, 81 | 1.11024 | 0.349870 | 0.945930 | 0.348436 |
| GAZE*MEM*HEMI | 1, 27 | 1.48574 | 0.233421 | 1.000000 | 0.233421 |
| GAZE*ROI*HEMI | 3, 81 | 0.17766 | 0.911248 | 0.685135 | 0.843294 |
| MEM*ROI*HEMI | 3, 81 | 0.35417 | 0.786236 | 0.895720 | 0.764275 |
| GAZE*MEM*ROI*HEMI | 3, 81 | 1.63482 | 0.187772 | 0.882338 | 0.193882 |
| **Alpha** | **df** | **F** | **p** | **H-F - Epsilon** | **H-F - Adj. p** |
| GAZE | 1, 27 | 1.83302 | 0.186996 | 1.000000 | 0.186996 |
| MEM | 1, 27 | 5.06462 | 0.032769 | 1.000000 | 0.032769 |
| ROI | 3, 81 | 23.55062 | 0.000000 | 0.802174 | 0.000000 |
| HEMI | 1, 27 | 13.07439 | 0.001211 | 1.000000 | 0.001211 |
| GAZE*MEM | 1, 27 | 1.61269 | 0.214950 | 1.000000 | 0.214950 |
| GAZE*ROI | 3, 81 | 1.48718 | 0.224225 | 0.984940 | 0.224794 |
| MEM*ROI | 3, 81 | 1.05910 | 0.371179 | 0.867080 | 0.365665 |

| | | | | | |
|---|---|---|---|---|---|
| GAZE*HEMI | 1, 27 | 0.02697 | 0.870782 | 1.000000 | 0.870782 |
| MEM*HEMI | 1, 27 | 2.34936 | 0.136969 | 1.000000 | 0.136969 |
| ROI*HEMI | 3, 81 | 2.84857 | 0.042554 | 0.832595 | 0.053287 |
| GAZE*MEM*ROI | 3, 81 | 1.22656 | 0.305460 | 0.915582 | 0.305192 |
| GAZE*MEM*HEMI | 1, 27 | 0.00449 | 0.947084 | 1.000000 | 0.947084 |
| GAZE*ROI*HEMI | 3, 81 | 2.60230 | 0.057593 | 0.900352 | 0.064318 |
| MEM*ROI*HEMI | 3, 81 | 1.43562 | 0.238476 | 0.631553 | 0.247360 |
| GAZE*MEM*ROI*HEMI | 3, 81 | 0.81824 | 0.487510 | 0.686850 | 0.449646 |

Section 3.4.2.3: The theta and alpha power after within-element saccades

| **Theta** | **df** | **F** | **p** | **H-F - Epsilon** | **H-F - Adj. p** |
|---|---|---|---|---|---|
| MEM | 1, 27 | 3.42955 | 0.075004 | 1.000000 | 0.075004 |
| ROI | 3, 81 | 31.07780 | 0.000000 | 0.860737 | 0.000000 |
| HEMI | 1, 27 | 3.31232 | 0.079867 | 1.000000 | 0.079867 |
| MEM*ROI | 3, 81 | 3.35742 | 0.022792 | 0.946770 | 0.025134 |
| MEM*HEMI | 1, 27 | 2.43335 | 0.130424 | 1.000000 | 0.130424 |
| ROI*HEMI | 3, 81 | 4.59780 | 0.005055 | 0.803769 | 0.009310 |
| MEM*ROI*HEMI | 3, 81 | 1.70443 | 0.172628 | 0.874536 | 0.179831 |
| **Alpha** | **df** | **F** | **p** | **H-F - Epsilon** | **H-F - Adj. p** |
| MEM | 1, 27 | 0.44376 | 0.510964 | 1.000000 | 0.510964 |
| ROI | 3, 81 | 13.52879 | 0.000000 | 0.793122 | 0.000004 |
| HEMI | 1, 27 | 6.47180 | 0.016995 | 1.000000 | 0.016995 |
| MEM*ROI | 3, 81 | 1.60122 | 0.195532 | 0.910576 | 0.199919 |
| MEM*HEMI | 1, 27 | 4.90865 | 0.035343 | 1.000000 | 0.035343 |
| ROI*HEMI | 3, 81 | 0.75253 | 0.524044 | 0.669494 | 0.476549 |
| MEM*ROI*HEMI | 3, 81 | 0.52645 | 0.665355 | 0.623931 | 0.582090 |
